# The Mechanism of Activation of Monomeric B-Raf V600E

**DOI:** 10.1101/2021.04.06.438646

**Authors:** Ryan C. Maloney, Mingzhen Zhang, Hyunbum Jang, Ruth Nussinov

**Affiliations:** Computational Structural Biology Section, Frederick National Laboratory for Cancer Research in the Laboratory of Cancer Immunometabolism, National Cancer Institute, Frederick, MD 21702, U.S.A; Department of Human Molecular Genetics and Biochemistry, Sackler School of Medicine, Tel Aviv University, Tel Aviv 69978, Israel

**Keywords:** B-Raf, oncogene, kinase, protein structure, activation mechanism

## Abstract

Oncogenic mutations in the serine/threonine kinase B-Raf, particularly the V600E mutation, are frequent in cancer, making it a major drug target. Although much is known about B-Raf’s active and inactive states, questions remain about the mechanism by which the protein changes between these two states. Here, we utilize molecular dynamics to investigate both wild-type and V600E B-Raf to gain mechanistic insights into the impact of the Val to Glu mutation. The results show that the wild-type and mutant follow similar activation pathways involving an extension of the activation loop and an inward motion of the αC-helix. The V600E mutation, however, destabilizes the inactive state by disrupting hydrophobic interactions present in the wild-type structure while the active state is stabilized through the formation of a salt bridge between Glu600 and Lys507. Additionally, when the activation loop is extended, the αC-helix is able to move between an inward and outward orientation as long as the DFG motif adopts a specific orientation. In that orientation Phe595 rotates away from the αC-helix, allowing the formation of a salt bridge between Lys483 and Glu501. These mechanistic insights have implications for the development of new Raf inhibitors.

## 1. Introduction

The Ras/Raf/MEK/ERK signaling pathway, also called the mitogen activated protein kinase (MAPK) pathway, is an important route for controlling cell life cycle events.[1–3] Dysregulation of the MAPK pathway negatively has been linked to several diseases including cancer and RASopathies, such as LEOPARD and Noonan syndromes.[4, 5] B-Raf, a member of the Raf family of serine/threonine protein kinases, is subject to a particularly large number of oncogenic mutations, the most prevalent of which is B-Raf V600E. B-Raf mutations are found in more than 50% of melanomas, as well as being prevalent in thyroid, ovarian, and colorectal cancers.[6] Due to the frequency of B-Raf mutations in cancer and other diseases, much of the activation mechanism of B-Raf has already been elucidated.[2] Several small molecule inhibitors of B-Raf V600E have been developed, yet long term patient outlook remains poor. Additional insights into the mechanism whereby the Val-to-Glu mutation causes B-Raf activation could provide key details in designing new oncogenic drugs.

The Raf family of serine/threonine protein kinases, which is composed of A-Raf, B-Raf, and C-Raf (also called Raf-1), is a key component of the MAPK pathway.[7] The members of the Raf family share three conserved regions (CRs) (Figure 1a).[8] CR1 is involved in autoinhibition and membrane recruitment and contains two subdomains: the Ras binding domain (RBD) and the cysteine-rich domain (CRD). CR2 is a flexible linker between CR1 and CR3 and contains a serine/threonine rich region that can be phosphorylated and binds to 14-3-3 scaffolding proteins. CR3 is the kinase domain that, upon activation, can phosphorylate its substrate proteins MEK1 and MEK2. The kinase domain consists of two lobes connected by a short hinge region (Figure 1b).[9] The N-terminal lobe, the smaller of the two, contains five β-strands and an α-helix called the C helix (αC-helix). Between β1 and β2 is the phosphate loop (P-loop, residues 462-469) which stabilizes ATP phosphate groups. The C-terminal lobe contains six α-helices as well as the activation loop (A-loop, residues 593-623) which begins with the conserved DFG motif (residues 594-596) and ends with the APE motif (residues 621-623). Between these two lobes is the ATP binding pocket.

**Figure 1.**
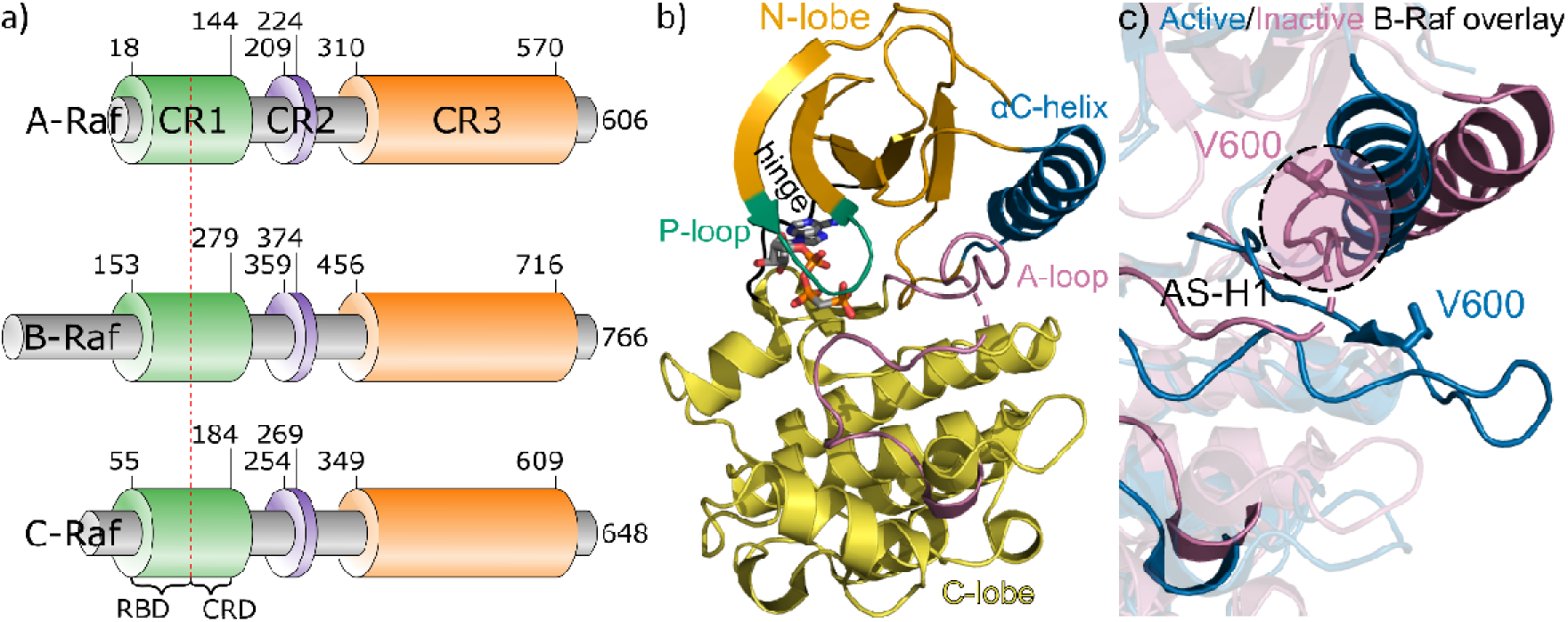
B-Raf structure and key features of the kinase domain. a) Conserved regions of Raf family proteins. b) B-Raf kinase domain crystal structures for inactive wild-type B-Raf (PDB ID: 6U2G). AMP-PCP (an ATP analog) is shown in the ATP binding pocket. c) Close up of A-loop and αC-helix for active B-Raf (PDB ID: 6UAN, blue) and inactive wild-type B-Raf (6U2G, pink). Missing residues are denoted by dashed lines.

Among the three members of the Raf family, B-Raf is subject to a particularly large number of oncogenic mutations. B-Raf mutations are found in more than 50% of melanomas, as well as being prevalent in thyroid, ovarian, and colorectal cancers.[6] These mutations are grouped into three classes.[10] Mutations involving B-Raf V600 residue belong to Class I. Class I mutants activate Raf through mimicking activation loop phosphorylation, causing B-Raf to adopt an active configuration.[11] Class II mutations are also kinase activated mutations independent from Ras. These mutations, however, occur on residues other than Val600, in particular residues of the AS-H1 inhibitory loop (residues 597-604), P-loop including Gly464 and Gly469, and activation segment involving Leu597 and Lys601.[10] The final class of B-Raf mutations, Class III mutations, also occur on residues other than V600; however, they are kinase impaired mutations. These Class III mutations require activation via Ras and heterodimerization with C-Raf.[12, 13] Nearly 90% of all B-Raf mutations are the Class I V600E mutation, and as such it is important to understand the activation mechanism of this mutation.[14, 15]

Wild-type B-Raf activation is highly regulated and involves a number of other proteins.[2] In the inactive state, wild-type B-Raf adopts a monomeric autoinhibited conformation in the cytosol that includes intramolecular interaction between the CR1 and CR3 regions.[16, 17] The autoinhibited B-Raf is often in a complex with MEK[18] and/or 14-3-3 scaffolding protein,[19] but notably not dimerized with a second Raf protein.[20] B-Raf is recruited to the membrane through a combination of interactions between the RBD with active Ras [21–23] and the CRD with the cell membrane.[24, 25] This releases B-Raf from the autoinhibited state and effectively increases the concentration of B-Raf near the cell membrane.[26] Upon recruitment to the membrane, B-Raf is activated through kinase domain dimerization, and two residues Thr599 and Ser602 in the A-loop are phosphorylated.[27–29] The phosphorylated residues destabilize the inactive state and stabilizes the active state. Once B-Raf adopts the active configuration, it is able to phosphorylate its substrate MEK1/2.[30, 31] The pseudo kinase, kinase suppressor of Ras (KSR) can act as a scaffolding protein to bring B-Raf and MEK into proximity with one another[32] and can also allosterically activate Raf.[33] The Hsp90/Cdc37 chaperone complex has also been implicated in A-Raf and C-Raf activity. Interestingly, inhibiting Hsp90 does not impact wild-type B-Raf function, but does decrease mutant B-Raf activity, including B-Raf V600E.[34] After B-Raf activates MEK, which in turn activates ERK, B-Raf returns to the inactive state through a series of phosphorylations of multiple sites on Raf. This results in breaking Raf/Raf and Raf/Ras dimerization.[35–38]

Switching from the inactive to the active conformation, B-Raf exhibits several characteristic structural changes.[8] In the inactive conformation, the αC-helix adopts an outward position and a collapsed A-loop with an inhibitory helix called AS-H1 (Figure 1c).[39] The DFG motif can also adopt an outward position in which the Phe595 sidechain flips outward to occupy the position normally occupied by the Asp594 sidechain, preventing ATP from entering the binding pocket.[40] In the active configuration, the αC-helix and DFG adopt an inward position and the A-loop extends, eliminating the inhibitory AS-H1 helix.[41] The inward position of the αC-helix causes the formation of a salt bridge between Glu501 of the αC-helix and Lys483 in the β3 strand.[4, 42, 43] The inward position of the DFG motif allows Asp594 to interact with Mg^2+^ bound to ATP, facilitating the transfer of the ATP γ-phosphate. Unlike wild-type B-Raf, which requires membrane recruitment and dimerization driven by active Ras for activation, B-Raf V600E adopts a catalytically competent structure independent of Ras.[39, 44] It has been proposed that the substitution of Val600 with the negatively charged Glu acts as a mimic of the two phosphorylated A-loop residues. It has also been suggested that a salt-bridge between Glu600 and Lys507 on the αC-helix in B-Raf V600E mutant is crucial for the formation of the active conformation.[39, 45] While many of these structural details of both wild-type and V600E B-Raf are known, additional details around the dynamic exchange between active and inactive conformations could provide novel targets for the development of inhibitors.

Previously it was believed that the Class I B-Raf V600E mutation created a constitutively active B-Raf monomer, independent of activation by Ras and Raf homo/hetero-dimerization.[46, 47] This theory was proposed based on experiments in which Arg509, an important residue for Raf dimerization, was mutated to His. The R509H mutation abrogates Raf dimerization, yet B-Raf V600E/R509H mutant cells still contained phosphorylated MEK and ERK, indicating monomeric Raf activity. Subsequent research, which included multiple mutations to the B-Raf dimer interface[11] and dimerization blocking peptides,[48] seemed to confirm that monomeric B-Raf V600E was able to phosphorylate MEK. Recent evidence disputes the theory that B-Raf V600E is constitutively active as a monomer.[49] Instead, it is argued, B-Raf V600E requires Raf homo/hetero-dimerization in order to phosphorylate MEK, just like wild-type B-Raf. This new perspective on B-Raf V600E is based on several factors. First, B-Raf V600E readily forms dimers or oligomers when expressed in cells.[11, 39, 50] Second, B-Raf V600E exhibits a larger dimer interface (DIF) compared to the wild-type B-Raf DIF, which allows B-Raf V600E dimer formation even in the presence of mutations to the dimer interface.[11] Third, the B-Raf V600E/ R509H/P622A mutant with two dimer impairing mutations (R509H in the dimer interface and P622A in the APE motif) exhibited greatly reduced dimer formation and completely abrogated MEK and ERK phosphorylation.[49] Finally, cells with B-Raf dimers in which one of the dimer pair is B-Raf V600E and the second a B-Raf mutant that is both kinase dead and unable to bind MEK showed inhibited cell growth and little to no ability to induce MEK or ERK phosphorylation. This indicates that B-Raf V600E needs a Raf partner able to hold MEK in order to stimulate cell growth, disputing the active monomeric B-Raf V600E theory.[49] Despite the growing evidence that B-Raf V600E signaling is dimer-dependent, probing the structure of the monomeric protein can have important implications in revealing the mechanism whereby the Val-to-Glu mutation disrupts autoinhibition and promotes dimerization. This is particularly important as research shifts to the development of dimer-breaking inhibitors.

B-Raf is a very attractive therapeutic target in the development of anticancer drugs due to the large number of oncogenic mutations in both B-Raf and its upstream activator Ras. The first-generation Raf inhibitors, including Vemurafenib (VEM) [51, 52] and Dabrafenib,[53] are ATP competitive inhibitors show promising early efficacy, but drug resistance quickly develops. [46, 47, 54–56] New inhibitors have been developed including pan-Raf inhibitors (such as LY3009120)[57] that bind to both protomers in Raf dimers, paradox breakers that selectively bind to B-Raf dimers, and allosteric inhibitors that target other sites apart from the ATP binding pocket.[58, 59] Despite this progress, long term survival rates remain low and additional treatment options are necessary.[60]

We perform molecular dynamics (MD) simulations on B-Raf wild-type and V600E mutant monomers. Unlike in previous work which simulated either B-Raf bound to inhibitors or with an empty ATP binding pocket, we have included ATP in our structures. In addition, we include models that contain phosphorylated A-loop with pThr599/pSer602 to replicate active wild-type B-Raf more closely. Results show that even with the V600E mutation, the AS-H1 helix is still present, however these residues do show greater fluctuation than in the wild-type simulations, indicating that the mutation has destabilized this inhibitory helix. In addition, simulations show that when the A-loop is extended, the αC-helix can move between the inward and outward positions if the DFG motif remains in an active configuration. These results aid in our understanding of the activation mechanism of B-Raf V600E. We provide detailed analysis linking the motion of the αC-helix to the orientation of the DFG motif. Importantly, these results indicate that the B-Raf V600E mutation results in an activation mechanism that mimics the mechanism of wild-type B-Raf.

### 2. Materials and Methods

Initial coordinates for the B-Raf kinase domain (residues 449 to 720, Figure S1) were adopted from the crystal structures (PDB IDs: 3OG7 [V600E active conformation], 4MNE [WT active conformation in complex with MEK], 4MNF [V600E active conformation], 4XV2 [V600E inactive conformation], 5ITA [WT intermediate conformation], 6U2G [WT inactive conformation in complex with MEK], 6U2H [WT active conformation in complex with 14-3-3]). Monomeric B-Raf structures were modeled by selecting the best resolved protomer in the crystal structures of Raf dimers, and any other proteins in the crystal structures were removed. Inhibitors present in the crystal structure were replaced with ATP and Mg^2+^. The initial configurations are shown in Figure S2 and structural details of these configurations are summarized in Table 1. To verify the binding between ATP/Mg^2+^ and B-Raf, we compare the interactions of our generated structures to the binding between B-Raf and AMP-PCP, an analog of ATP (PDB ID: 6U2G) in Table S1. The network analysis with Maestro[61] on the initial configurations illustrates that the binding modes between the ATP/Mg^2+^ molecules and B-Raf are similar for all systems (Figure S3). We focused on initial conditions with the DFG-in configuration, as the DFG-out configuration would cause the Phe595 to overlap with the ATP phosphates. From there, we selected Raf structures in which the αC-helix exhibited either an inward, outward, or intermediate orientation. The shorter loop segments without coordinates in the ATP binding P-loop and other loops in the C-lobe were modeled using the CHARMM program.[62] Any mutated residues present, except for residue 600, were changed back to the wild-type sequence (see Table S2). Residue 600 was either kept as Val to obtain the B-Raf wild-type (configurations *j*-*p*) or mutated to Glu obtain B-Raf V600E (configurations *a*-*i*). In addition, configurations *n*-*p* contain phosphorylated A-loop with pThr599/pSer602. The A-loop was modeled to have either an “extended” or “collapsed” configuration. For the extended A-loop, the longer loop segments without coordinates were constructed with SWISS-MODEL,[63] adopting available loop conformations in a database from PDB. A series of minimization and dynamic cycles for the constructed loops with constrains on the defined regions were performed to optimize the unstructured loops. Structures with a collapsed A-loop, apart from configuration *h*, exhibit the AS-H1 inhibitory helix in the initial A-loop structure. For configuration *h* the inhibitory helix is not present, but A-loop residues are positioned near the main body of the protein.

**Table 1.** Initial Configurations of the simulated systems of monomeric B-Raf. The distance between Lys483 and Glu501 is measured between the β-carbon atoms and is calculated after modeling missing residues and performing energy minimization steps. *R_g_* is the radius of gyration.

| | configuration | PDB | K483 - E501<br>Distance (Å) | αC -helix | A-loop | AS-H1 | A-loop<br>Rg (Å) |
| --- | --- | --- | --- | --- | --- | --- | --- |
| V600E: | a | 4MNE | 7.81 | in | extended | no | 0.86 |
|  | b | 4MNE | 9.07 | in | extended | no | 0.88 |
|  | c | 6U2H | 7.81 | in | extended | no | 0.82 |
|  | d | 3OG7 | 8.32 | in | extended | no | 0.76 |
|  | e | 4MNF | 7.89 | in | extended | no | 0.87 |
|  | f | 5ITA | 10.16 | intermediate | extended | no | 0.85 |
|  | g | 5ITA | 10.75 | intermediate | collapsed | yes | 0.62 |
|  | h | 4XV2 | 12.26 | out | collapsed | no | 0.68 |
|  | i | 6U2G | 13.03 | out | collapsed | yes | 0.64 |
| wild-type: | j | 4MNE | 8.03 | in | extended | no | 0.86 |
|  | k | 6U2H | 7.37 | in | extended | no | 0.79 |
|  | l | 5ITA | 11.12 | intermediate | collapsed | yes | 0.67 |
|  | m | 6U2G | 14.16 | out | collapsed | yes | 0.60 |
| pT599/pS602<br>wild-type: | n | 4MNE | 7.30 | in | extended | no | 0.77 |
|  | o | 5ITA | 10.88 | intermediate | collapsed | yes | 0.67 |
|  | p | 6U2G | 14.80 | out | collapsed | yes | 0.63 |

The B-Raf monomers with ATP and Mg^2+^ were placed in a periodic box of ~100 × 100 × 100 Å^3^ and the explicit TIP3 water model was used to solvate the system. Water molecules within 2.6 Å of B-Raf were removed to prevent atom collapse. Sodium (Na^+^) and chlorine (Cl^−^) ions were added to generate a final ionic strength of ~100 mM and neutralize the system. During the preequilibration stage, a series of minimization cycles were performed for the solvents including ions with the harmonically restrained protein backbone until the solvent reached 310 K. Following the energy minimization steps, a series of dynamic cycles were performed by gradually releasing the harmonic constraints on the protein backbone with the long-range electrostatics calculation using the particle mesh Ewald (PME) method. The production runs of all-atom MD simulations were conducted using the NAMD package (version 2.12)[64] with the updated CHARMM[62] all-atom force field (version 36m).[65, 66] All simulations closely followed the same protocol as in our previous studies.[25, 67–83] In the productions runs, the Langevin thermostat maintained the constant temperature at 310 K and the Nosé-Hoover Langevin piston pressure control sustained the pressure at 1.01325 bar (1 atm) with the NPT condition. A 2 fs timestep was used for 5 × 10^8^ steps for all simulations. A total of 16 μs simulation were performed for 16 systems, each with 1 μs simulation time. Trajectory information was collected every 5 × 10^4^ steps (100 ps).

## 3 Results

### 3.1 Wild-Type B-Raf Kinase Domain

First, we look at features present in the active and inactive conformations of wild-type B-Raf. To get a broad picture of the movement of B-Raf, Figure 2a and b show the root mean square fluctuation (RMSF) for wild-type and pThr599/pSer602 B-Raf simulations (Results for all simulations provided in Figure S4). In general, the simulations show a flexible region at the P-loop, the loop connecting β3 and the αC-helix (including the first few residues of the αC-helix), the residues of β4 through β5, the A-loop, the loop N-terminal of the αF-helix, and the residues around the αG-helix. These regions are highlighted in red in the snapshots in Figure 2c for wild-type B-Raf with an initial inward (configurations *j* in Table 1), intermediate (*l*), or outward αC-helix (*m*), which are colored according to *B*-factor values. We compare these RMSF and *B*-factor results with the *B*-factor data available from crystal structures used to create our initial configurations (Figure S5). For the most part, the regions were found to have high RMSF values are the same regions with high *B*-factor values in the deposited crystal structures. Indeed, the P-loop, β3 to αC loop, A-loop, and αF-helix N-terminal loop residues are all lacking density in at least one of the crystal structures. There were two key differences between the simulation data and the crystal structures. First, most crystal structures (except for 6U2H) had lower *B*-factor values around the αG-helix, particularly those crystal structures of Raf/MEK dimers. The αG-helix is important in B-Raf/MEK binding,[84] so we would expect these residues to be more flexible in monomers. Indeed, previous simulations of monomeric B-Raf with an empty ligand binding pocket have found a possible active-inactive transition state in which this helix is partially unfolded.[45] Second, most of the crystal structures display pronounced peaks near residues 545, 585, and (for simulations that do not contain other proteins; 3OG7, 4MNF, 4XV2, 5ITA) the residues between 680 and 690. The first of these peaks corresponds to a loop between αD and αE. The second peak corresponds to a loop between β7 and β8, just before the start of the A-loop. The third peak corresponds to the loop at the N-terminal of the αH-helix. These regions, which are indicated in the center panel of Figure 2c, are near each other on the surface of the protein. The proximity of these loops could indicate a cooperative stabilization in the simulation.

**Figure 2.**
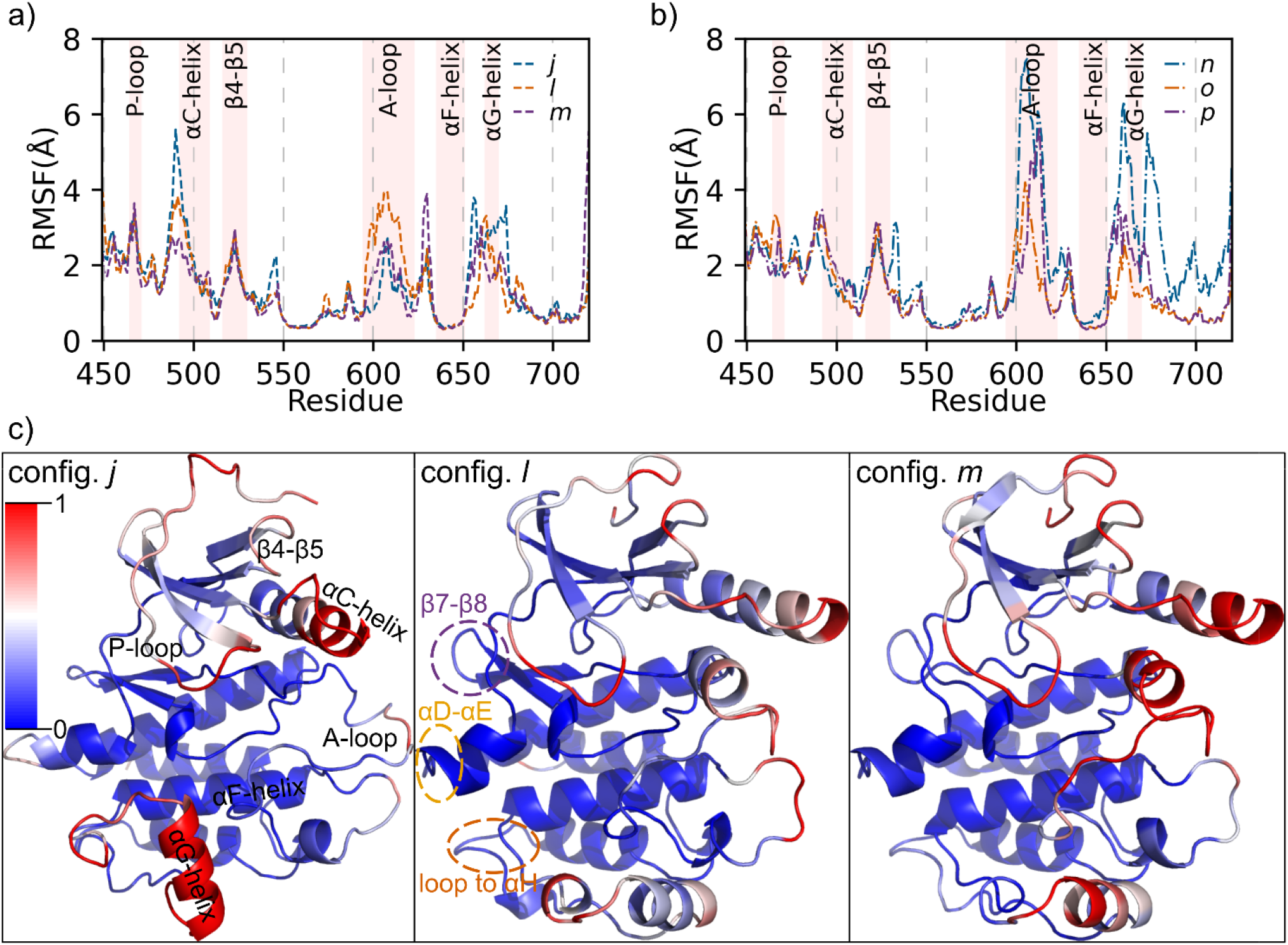
B-Raf exhibits peaks in residue fluctuation in the P-loop, residues before the αC-helix, A-loop, and around the αG-helix. a) RMSF for residues in wild-type B-Raf configurations *j* (blue), *l* (orange), and *m* (purple). b) RMSF for residues in pThr599/pSer602 B-Raf configurations *n* (blue), *o* (orange), and *p* (purple). c) Snapshot of configuration *j*, *l*, and *m* colored by normalized *B*-factor value from maximum *B*-factor (red) to minimum *B*-factor (blue).

Next, we look more specifically at the impact that αC-helix position and activation loop phosphorylation have on RMSF. In Figure 2a, we see that the β3-αC loop is most flexible for configuration *j* and the A-loop is most flexible for configuration *l*. For configuration *j*, an active form of monomeric wild-type B-Raf (with an inward αC-helix) is expected to be less stable than the inactive form (with an outward αC-helix), thus the high RMSF values in this loop could be due to movement of the αC-helix. For configuration *l*, the intermediate position of the αC-helix could cause destabilization of the A-loop residues. In Figure 2b, both configuration *n* and *p* have large RMSF values for the A-loop, and configuration *n* also displays peaks around the αG-helix. The high RMSF values of the A-loop, particularly when compared to the unphosphorylated systems, could indicate that the phosphate motif destabilizes the activation loop. These regions are investigated in more detail below to identify the differences in the active and inactive wild-type B-Raf structures.

A widely reported hallmark of active kinases is an “inward” position of the αC-helix. This inward position allows formation of a salt bridge between Lys483 of the β3 strand and Glu501 of the αC-helix of B-Raf. We plot the distance between the β carbon (CB) atoms on Lys483 and Glu501 in Figure 3a and b for the wild-type and pThr599/pSer602 simulations, respectively. The results from all simulations are in Figure S6. This distance indicates if the Lys483-Glu501 salt bridge is present, which gives an indication of the αC-helix position throughout the simulations. Simulations that began with an inward αC-helix (configurations *j*, *n*) exhibited a mostly stable salt-bridge that had a distance of ~8.2 - 8.75 Å between the CB atoms. There is, however, a region of ~700 ns during which the salt bridge of configuration *j* repeatedly broke and reformed. We can see from the snapshots in Figure 3c that when the salt bridge is broken the αC-helix moves up and outward and when the salt bridge is present, the αC-helix moves down and inward. This inward and outward motion of the αC-helix could account for the large RMSF values of configuration *j* in this region. This is supported by the lack of inward and outward motion seen for configuration *n*, which had a corresponding lower RMSF in this region. The salt bridge gradually moved apart over the course of the simulations for configurations that began with an intermediate αC-helix and collapsed A-loop (configurations *l*, *o*). In configuration *l*, the Lys483 and Glu501 residues begin at a distance of ~12 Å and move closer together for the first ~300 ns, after which these residues move gradually apart for the remainder of the simulation, to a distance of ~13 Å. In configuration *o*, the salt bridge begins at a distance just under 10 Å, and moves slightly closer for the first ~100 ns and then moves gradually apart for the remainder of the simulation, to a distance of ~11 Å. The simulations that began with an outward αC-helix (configurations *m*, *p*) maintained this orientation throughout the course of the simulation, with the distance between Lys483 and Glu501 of ~15 Å. The position of the αC-helix can also be measured by calculating the distance between the amide nitrogen (NZ) in the sidechain of Lys483 and the carboxylate carbon (CD) in the sidechain of Glu501. The results of these calculations are shown in Figure S7a. Also presented in Figure S7b is the center of mass distance between the αC-helix in the N-lobe and αE-helix in the C-lobe.[85] These data follow the same trends as shown in Figure 3 and show that the formation of the salt bridge between Lys483 and Glu501 is tied to the inward motion of the αC-helix.

**Figure 3.**
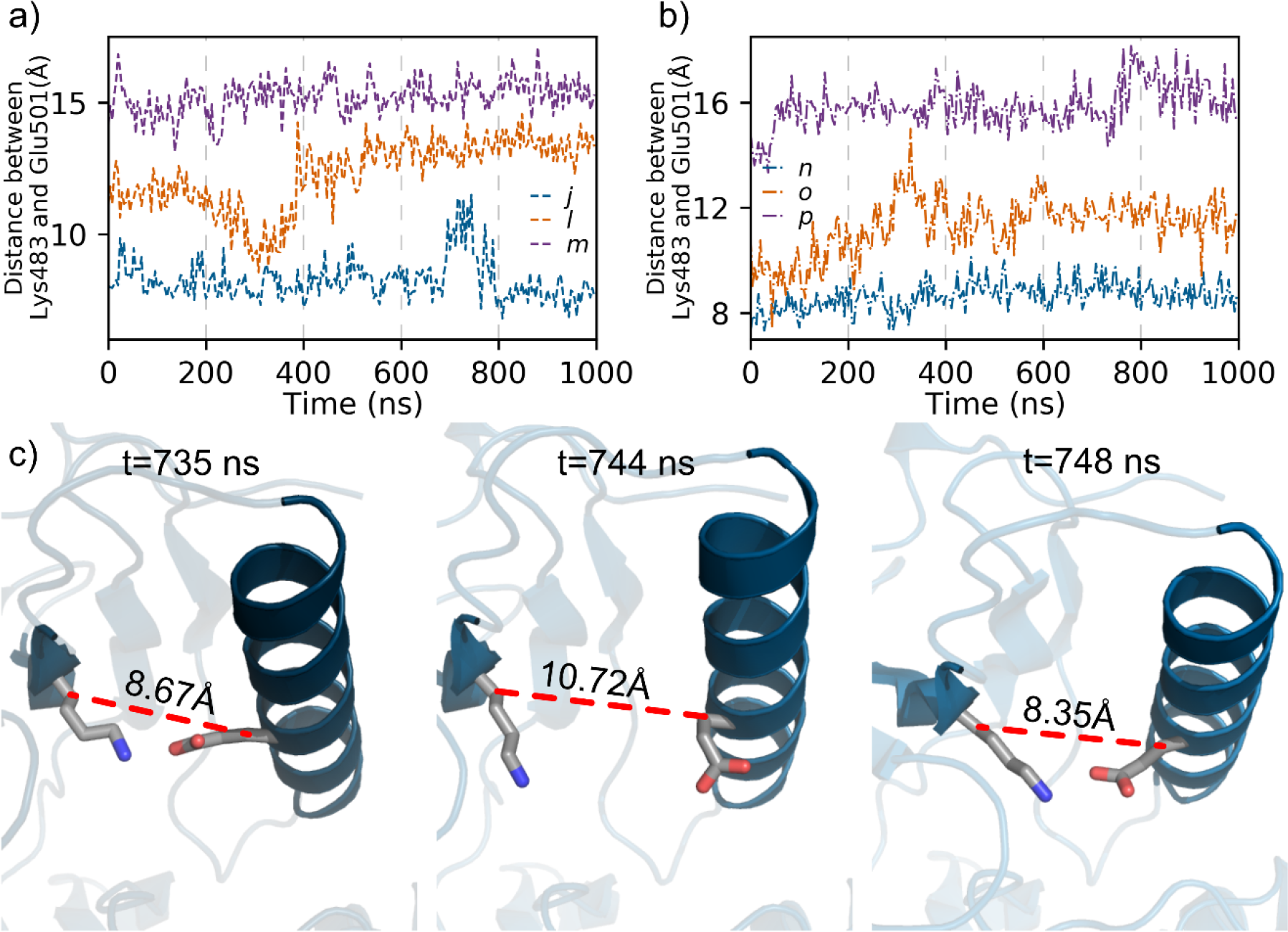
Wild-type B-Raf αC-helix position. a) Distance between Lys483 and Glu501 over the course of the simulations for wild-type B-Raf configurations *j* (blue), *l* (orange), and *m* (purple). Distance between Lys483 and Glu501 over the course of the simulations for pThr599/pSer602 B-Raf configurations *n* (blue), *o* (orange), and *p* (purple). c) Snapshots representing the salt bridge interaction between Lys483 and Glu501 at selected times of *t* = 735, 744, and 748 ns for configuration *j*.

A second important feature that distinguishes active and inactive B-Raf is an extended activation loop for the active kinase. Although it is not expected to be the most stable configuration, monomeric wild-type B-Raf simulations that started with an extended A-loop maintained it throughout the simulation. Configuration *j* shows that the extended A-loop may be stabilized through hydrophobic interactions between Val600, Val504, Ile572, and Ala598 along with hydrogen bonds between β6 and β9 (Figure 4a).[86] Configuration *m* shows that the collapsed A-loop with inhibitory AS-H1 helix is stabilized through formation of a hydrophobic pocket that includes Val600 along with residues Leu485, Val487, Phe498, Val502, Phe516, Ile527, and Leu597 (Figure 4b). Configuration *l* shows that the extended A-loop with the inhibitory AS-H1 helix does not form these stabilizing interactions: Val600 is not in a hydrophobic pocket and β9 is not positioned to create hydrogen bonds with β6 (Figure 4c). This could explain why the RMSF values for the A-loop of configuration *l* is larger than those of configurations *j* and *m*. With the phosphorylation of residues Thr599 and Ser602, charge interactions can explain the RMSF values of the A-loop in configurations *n*-*p*. In Figure 4d, we see the same hydrophobic and hydrogen bonding interactions in configuration *n* (faded yellow and black residues) that was present in configuration *j*. However, after residue 600, the A-loop of configuration *n* extends further into the solvent than the A-loop of configuration *j* did. This allows pSer602 to interact with Arg603, water, salt ions in the solution, and places it near Lys507. These charged interactions could cause this region of the A-loop to fluctuate. The pThr599, on the other hand, is more internal in the protein and interacts with Lys601 and Arg575, the HRD arginine residue. The position of pThr599 would minimize its interactions with the solvent and could account for the low RMSF values in the A-loop up to residue 600 in configuration *n*. Figure 4e shows that in configuration *p*, Val600 again sits in a hydrophobic pocket seen in configuration *m*, which would suggest the RMSF values of the A-loop would be low. However, the phosphorylated residues, which are located on the inhibitory AS-H1 helix, are situated on the surface of the protein and near the negatively charged Glu501. These repulsive interactions, plus solvent interactions, could account for the high RMSF values of the activation loop of this configuration. Finally, configurations *l* and *o* had similar RMSF values in the A-loop. Figure 4f shows that configurations *o* had very few hydrophobic interactions around Val600, like the unphosphorylated configuration *l*. For the phosphorylated residues in configuration *o*, pSer602 is facing toward the internal part of the protein and interacts with Arg575, while pThr599 is facing toward the solvent and is situated near Arg603. The charge-charge interactions between the phosphorylated residues and their neighboring residues do not seem to add any stability to the A-loop. Interestingly, the relative positions of the phosphorylated residues of configuration *o* are opposite of those in the active configuration *n,* which indicates substantial rearrangement of the A-loop is necessary for activation.

**Figure 4.**
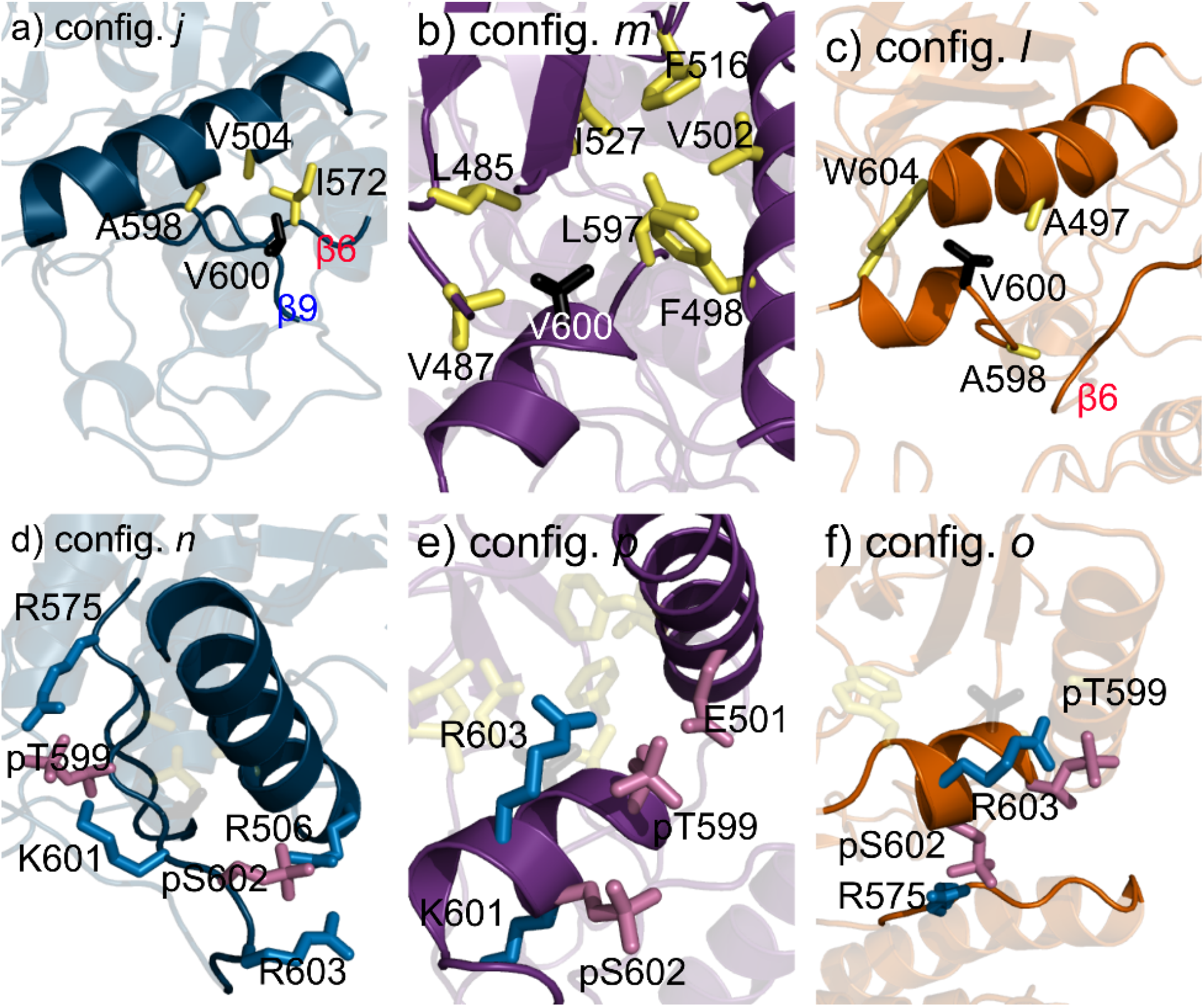
Hydrophobic interactions between V600 and neighboring residues stabilize the extended or collapsed A-loop. a) Hydrophobic residues (yellow) around Val600 (black) and hydrogen bonding between β6 (red) and β9 (blue) stabilize the extended A-loop in configuration *j*. b) Hydrophobic residues form a pocket that contains Val600 and stabilizes the collapsed A-loop configuration in configuration *m*. c) The extended A-loop with inhibitory helix of configuration *l* lacks the large hydrophobic pocket of configuration *m*, and the hydrogen bonding β-sheet interactions of configuration *j*. d) Positively charged residues (blue) near pThr599/pSer602 (pink) in configuration *n*. e) pThr599/pSer602 (pink) residues near negatively charged residue Glu501 and in configuration *p*. f) pThr599/pSer602 residues in configuration *o* are in opposite positions relative to those of configuration *n*.

### 3.2 Active B-Raf V600E

The B-Raf V600E simulations that began with an inward αC-helix behaved much the same as an inward αC-helix wild-type B-Raf. Superimposed snapshots in Figure 5a of the active wild-type (configuration *j*) and mutant B-Raf (configurations *a*-*d*) show that they largely share the same structural features. In these snapshots, the residues of the A-loop after Ser605 and the residues around the αG-helix (Gln653 to Val681) show the largest differences in their positions between the wild-type and mutant configurations. These residues also display the greatest values in the RMSF as shown in Figure 5b. RMSF results also show that active B-Raf V600E generally displays larger values in the activation loop than the active wild-type B-Raf, but similar values to those seen in pThr599/pSer602. This indicates that the charged Glu residue introduced in the V600E mutation behaves in a similar manner to the charged phosphorylated residues in wild-type B-Raf. The Lys483-Glu501 salt bridge is stable through the course of the simulations for configurations *a*-*d*. In configuration *e* the salt bridge broke and reformed throughout the final 700 ns of the simulation (see Figure 5c and Figures S6, S7), spending most of this time in an outward orientation with rapid swings inward and then back out. This is similar to the breaking and reforming of the salt bridge that was seen in wild-type B-Raf configuration *j* (Figure 3a). Unlike configuration *j*, which ended the simulation with an inward αC-helix, the B-Raf V600E configuration *e* showed an outward shift in the αC-helix along with the breaking of the salt bridge, as can be seen in Figure S8.

**Figure 5.**
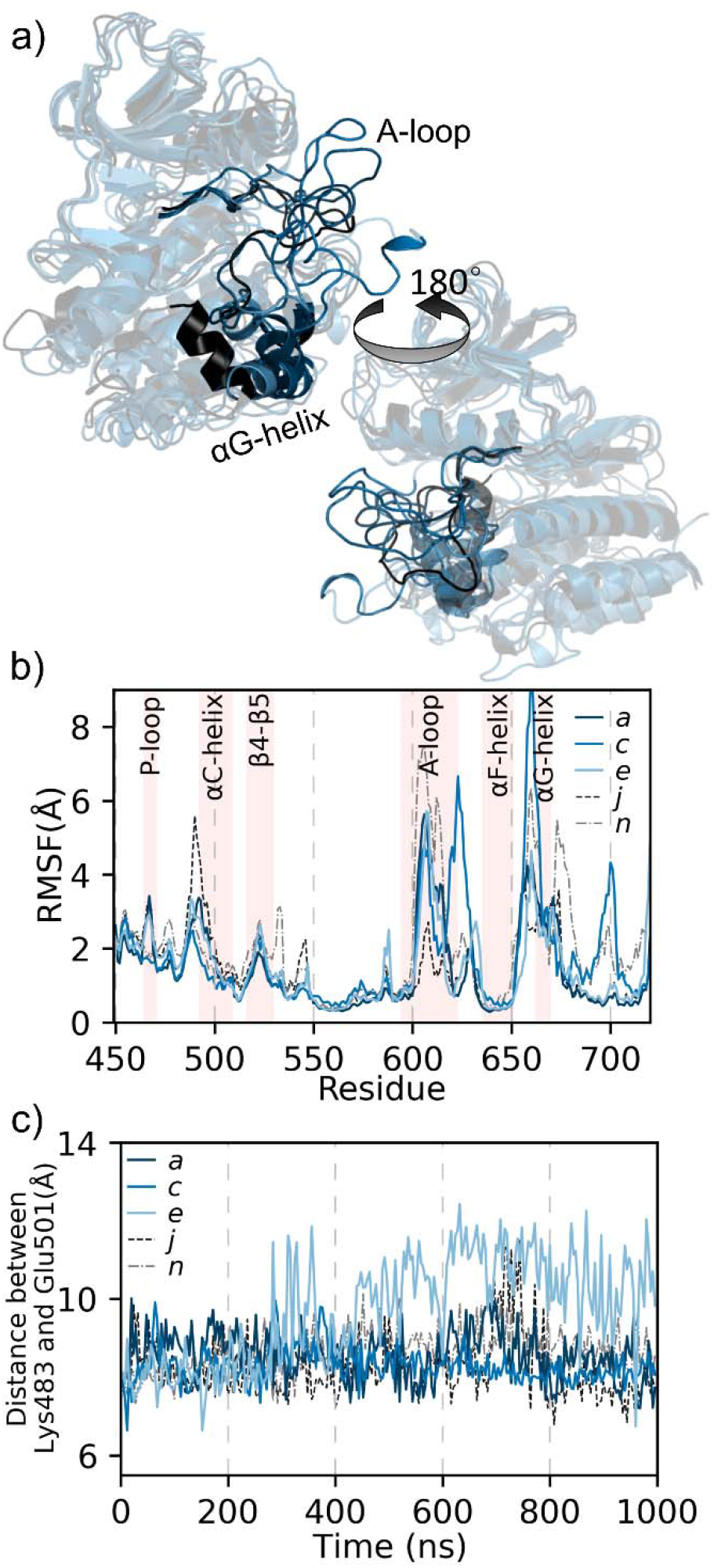
Active B-Raf V600E exhibits many of the same features as active wild-type B-Raf. a) Snapshots of active wild-type B-Raf (configuration *j*, black) and B-Raf V600E (configurations *a d*, dark blue to light blue). The αG-helix and activation loop are indicated. b) RMSF for kinase domain residues for B-Raf V600E configurations *a* (dark blue), *c* (medium blue), and *e* (light blue) that began with an inward αC-helix. c) Distance between Lys483 and Glu501 over the course of the simulations for B-Raf V600E configurations that began with an inward αC-helix: *a* (dark blue), *c* (medium blue), and *e* (light blue). A comparable wild-type B-Raf, configuration *j*, and pThre599/pSer602, configuration *n*, are shown in the black dashed and gray dash dot line, respectively.

In the active wild-type B-Raf, Val600 formed hydrophobic interactions with two nearby residues while the A-loop was extended. In B-Raf V600E, it has been proposed that the Glu600 forms a salt bridge with Lys507. Enhanced-sampling simulations have found that this salt bridge is not stable and have instead proposed a possible salt bridge between Glu600 and Arg603.[45] We plot the distances between the Glu600 carboxylate carbon (CD) and the Lys507/Arg603 amide nitrogen (NZ) in Figure 6a and 6b, respectively. This shows that the Lys507-Glu600 salt bridge remains relatively stable through most of the simulation, but it does break and quickly reform periodically. The salt-bridge between Arg603 and Glu600 is much less stable, though may play a secondary role in stabilizing the extended A-loop.

**Figure 6.**
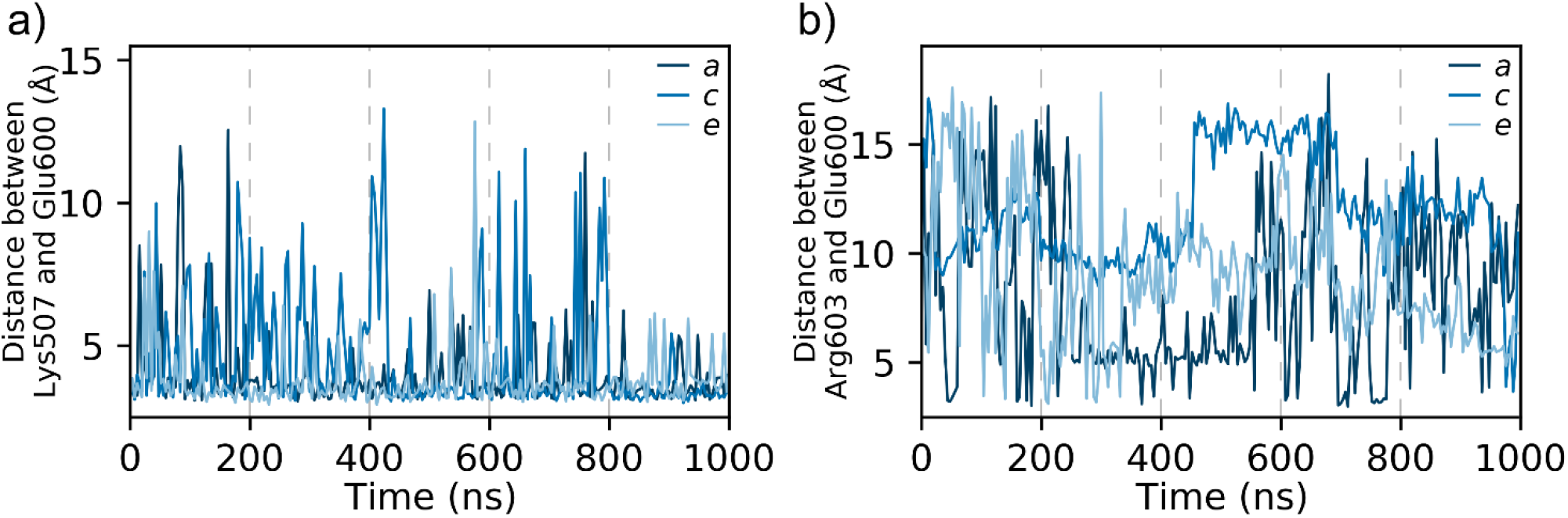
The salt bridge between Glu600 and Lys507 stabilizes the extended A-loop in active B-Raf V600E. Distance between Glu600 and a) Lys507 and b) Arg603 for configurations that began with an inward αC-helix: *a* (dark blue), *c* (medium blue), and *e* (light blue).

### 3.3 Inactive B-Raf V600E

The simulations that begin with intermediate or outward oriented αC-helix display distinct behavior from their wild-type counterparts. Although none of these simulations moved from the inactive to the active state, as other simulations have reported,[85] there are indications how this mutation may destabilize the inactive configuration. Figure 7a shows that the A-loop residues for the simulations that began with the αC-helix in an intermediate position (configurations *f* and *g*) had larger RMSF values than both the wild-type and pThr599/pSer602 (configurations *l* and *o*, respectively). Figure 7b shows that the A-loop residues for the simulations that began with the αC-helix in an outward position (configurations *h* and *i*) had larger RMSF values than wild-type B-Raf (configuration *m*), but closely matched the RMSF values of pThr599/pSer602 B-Raf (configuration *p*). This result supports the hypothesis that the V600E mutation mimics the activation loop phosphorylation in wild-type B-Raf. One mechanism where the Val-to-Glu mutation could induce an active conformation in B-Raf is through destabilizing the hydrophobic pocket occupied by Val600 in wild-type B-Raf. Figure 7c shows that the αC-helix behavior of the V600E mutant simulations with an intermediate αC-helix in the initial configuration (configurations *f* and *g*) is much more similar to that of pThr599/pSer602 B-Raf (configuration *o*) than to that of the wild-type B-Raf (configuration *l*). In these V600E mutant systems, like the pThr599/pSer602 system, the Lys483 and Glu501 residues remain ~10 Å apart and did not move away from each other, as was seen in the wild-type system. The reason that the V600E and pThr599/pSer602 systems remain closer may be due to repulsive interactions between Glu501 with either Glu600 or the phosphorylated residues. As shown in Figure 7d, these negatively charged residues are near one another, and the αC-helix in configurations *f*, *g*, and *o* are displaced further away from the activation loop than the helix in configuration *l*, which lacks a negatively charged residue around residue 600. By moving the αC-helix away from the A-loop, Glu501 is positioned closer to β3, which could aid in the key salt bridge formation and activation. Figure 7e shows that the configurations that began with an outward αC-helix maintained this orientation throughout the simulation.

**Figure 7.**
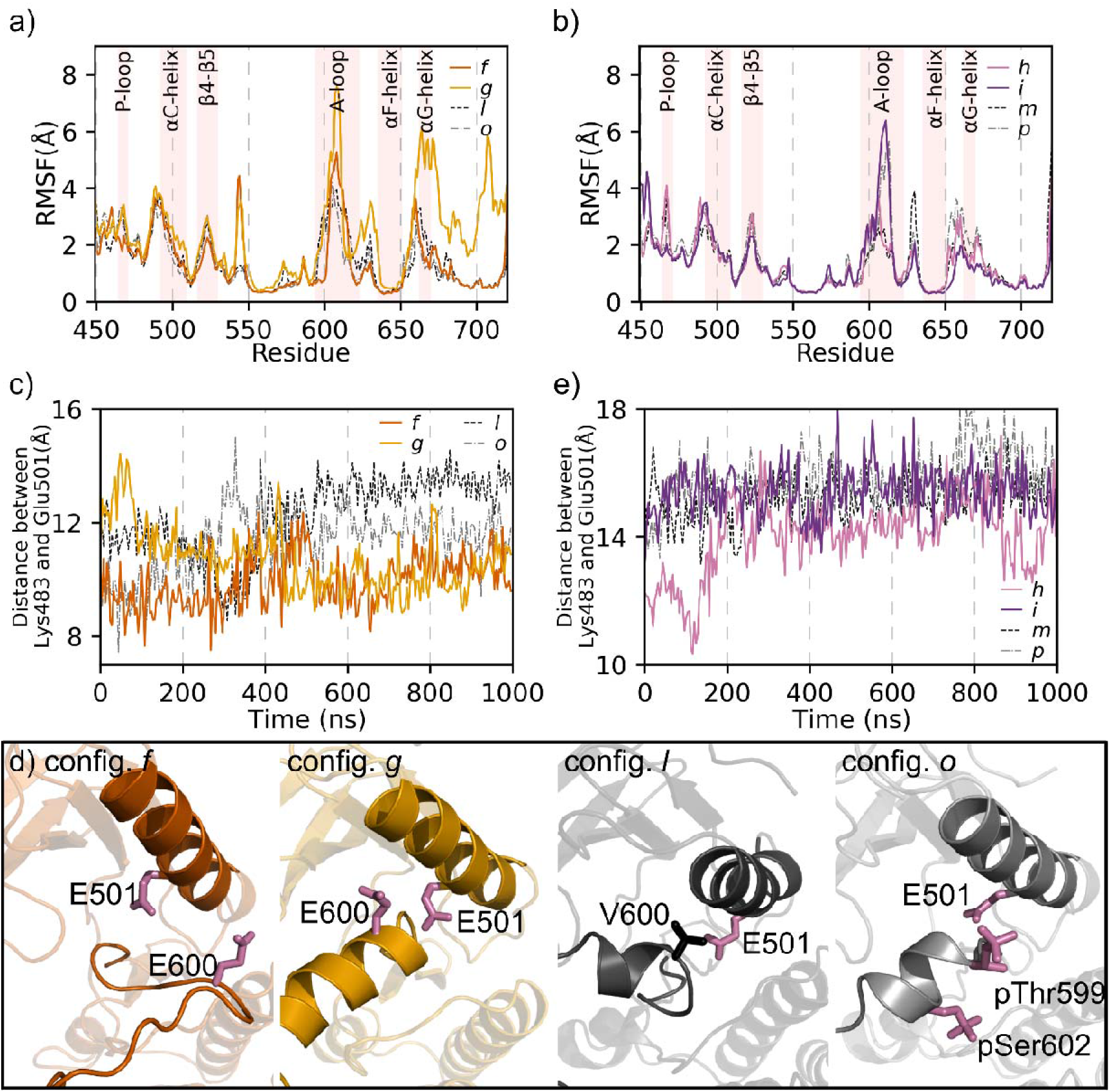
B-Raf V600E with intermediate and outward αC-helix do not change to an inward αC-helix. RMSF for kinase domain residues for B-Raf V600E configurations that began with an a) intermediate αC-helix, *f* (dark orange) and *g* (light orange), and b) outward αC-helix, *h* (light purple) and *i* (dark purple). A comparable wild-type and pThr599/pSer602 B-Raf system is shown in the black dashed and gray dash dot line, respectively, in each figure. c) Distance between Lys483 and Glu501 through the course of the simulation for B-Raf V600E configurations that began with an intermediate αC-helix. d) Snapshot of αC-helix position at the end of the simulation for configurations with an intermediate αC-helix. e) Distance between Lys483 and Glu501 through the course of the simulation for B-Raf V600E configurations that began with an outward αC-helix.

The inactive mutant systems also display different Glu600 salt bridge formation than was seen in active B-Raf V600E. Configuration *f*, which had an extended A-loop in the initial configuration, did show the formation of salt bridge between Lys507 and Glu600 (Figure 8a). The other inactive mutant systems, configurations *g*, *h*, and *i*, had collapsed A-loops and did not form this salt bridge. Instead, these inactive B-Raf V600E systems showed that the salt bridge between Glu600 and Arg603 (Figure 8b), was much more prevalent than was seen in the active B-Raf V600E systems (Figure 6b). The formation of this salt bridge could stabilize the collapsed A-loop with the inhibitory AS-H1 helix, even though the V600E mutation would disrupt the hydrophobic pocket occupied by V600, shown in Figure 4b.

**Figure 8.**
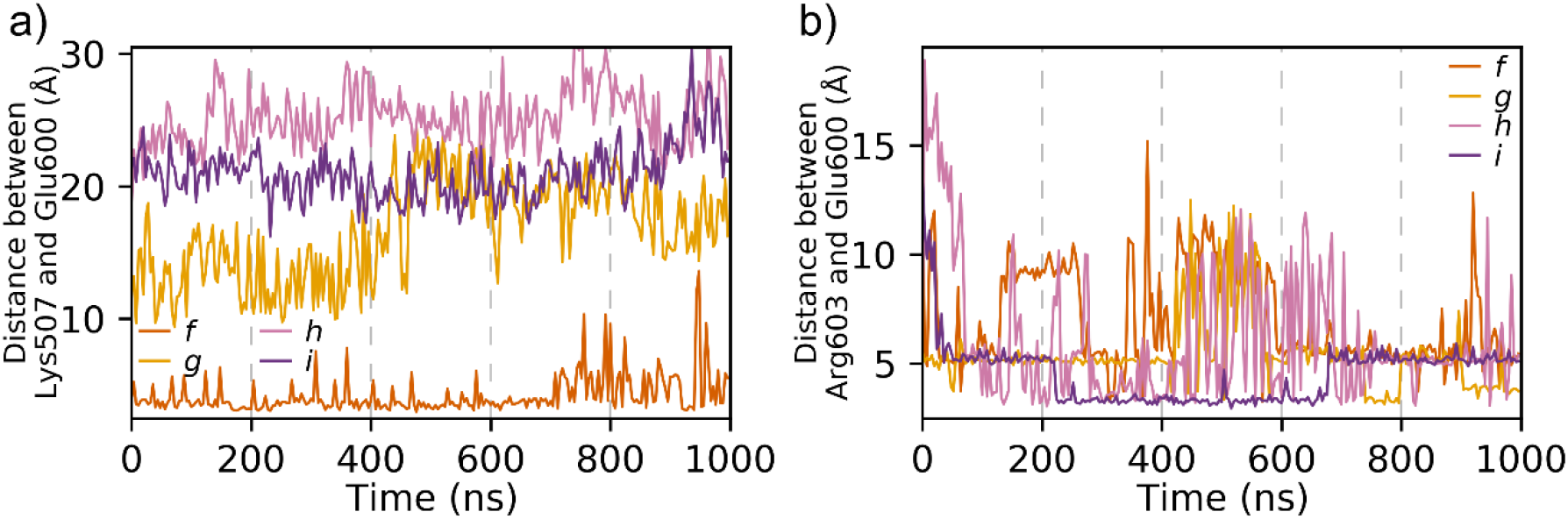
The salt bridge between Glu600 and Arg603 stabilizes the collapsed A-loop in inactive B-Raf V600E. Distance between Glu600 and a) Lys507 and b) Arg603 for B-Raf V600E configurations that began with an intermediate or outward αC-helix: *f* (dark orange), *g* (light orange), *h* (light purple), and *i* (dark purple).

### 3.4 Link between activation loop and αC-helix position

To understand the differences in the activation loop behavior of B-Raf, we look at the DFG motif (residues 594-596), which is located at the beginning of the A-loop. Inhibitors are known to bind to B-Raf with the Phe residue of the DFG motif in either an in or out orientation. In the out orientation, the phenylalanine ring occupies the ATP binding pocket and is thus incompatible with our monomeric B-Raf with ATP simulations. Indeed, when we measure the position of this residue, we find that it maintains an inward orientation throughout the course of the simulation. Of more interest is the orientation of the dihedral angles around the XDFG motif (DFG motif plus the preceding Gly593 in B-Raf). It has been proposed that kinases can be classified as active or inactive based on the backbone dihedrals □ and ψ of the residues, Gly593, Asp594, and Phe595, along with the first sidechain dihedral, χ_1_, of Phe595.[87] Specifically, they found that all catalytically primed structures exhibit a “BLAminus” conformation, that is the backbone dihedrals of the X residue (Gly593) occupies the β (B) region of Ramachandran map, Asp594 occupies the left (L) region, and Phe595 occupies the α (A) region. Finally, the sidechain of Phe595 adopts a χ_1_ gauche-minus (−60°) rotamer. The BLAminus conformation is the only one that properly positioned the DFG motif and A-loop to simultaneously bind ATP, magnesium ion, and substrate. Figure S9 shows how the XDFG dihedral angles evolved over the course of each simulation. Simulations that began with an inward αC-helix and extended A-loop all displayed the BLAminus XDFG dihedral orientation throughout the course of the simulation (Figure S9, configurations *a*-*e* for B-Raf V600E, *j*-*k* for wild-type B-Raf, and *n* for pThr599/pSer602 B-Raf). An example of this can be seen for configuration *j* in Figure 9a. In this simulation, B-Raf V600E maintains the BLA configuration for the backbone dihedrals, even during the time period when the αC-helix switches repeatedly between the inward and outward orientation (Figure 3a). The Phe595 χ_1_ dihedral oscillates between −90° and −50° throughout the simulation (Figure 9b). This fluctuation can be expected from the nature of molecular dynamics simulation, but importantly the χ_1_ angle never switches to trans (180°) or gauche-plus (60°). A snapshot of the DFG region for this configuration is shown Figure 9c. Therefore, we can conclude that this configuration adopts the BLAminus orientation and is an active kinase. The inward αC-helix B-Raf simulations (configurations *a*-*e*, *k*, *n*) also exhibit the BLAminus XDFG orientation, even configuration *e* when the salt bridge was breaking and reforming (Figures S6 and S7).

**Figure 9.**
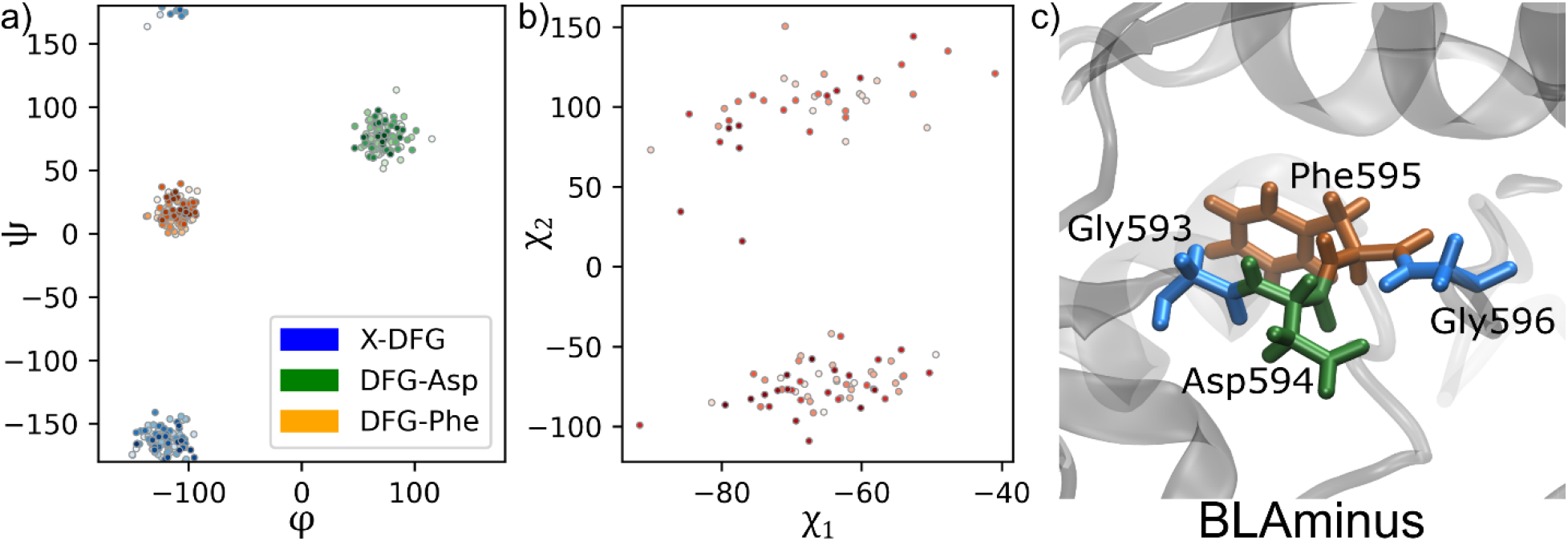
Active B-Raf adopts a BLAminus XDFG orientation. a) Ramachandran graph, b) sidechain dihedral angles, and c) representative snapshot of the residues ^593^GDFG^596^ for configuration *j*. Lightly colored symbols represent data from early in the simulation, darkly colored symbols are from the end of the simulation.

One of the B-Raf V600E simulations with an intermediate αC-helix, configuration *f*, had an initial XDFG orientation of BLAminus but changed to BLAtrans in the final ~300 ns of the simulation (Figure 10a, b). The change in the Phe595 χ_1_ gauche rotamer from minus to trans moves the phenylalanine sidechain into the pocket between the αC-helix and β3 (Figure 10c). Figure 7c showed that this configuration did have an inward orientation of the αC-helix through much of the first ~700 ns of the simulation, but not for the final 300 ns. This can be seen even more clearly by plotting the distance between the amide nitrogen (NZ) in the sidechain of Lys483 and the carboxylate carbon (CD) in the sidechain of Glu501 (Figure 10d). This shows that after the Phe595 χ_1_ gauche rotamer becomes trans, the Lys483-Glu501 salt-bridge can no longer form and highlights the importance of the positioning of this residue on kinase activation.[88]

**Figure 10.**
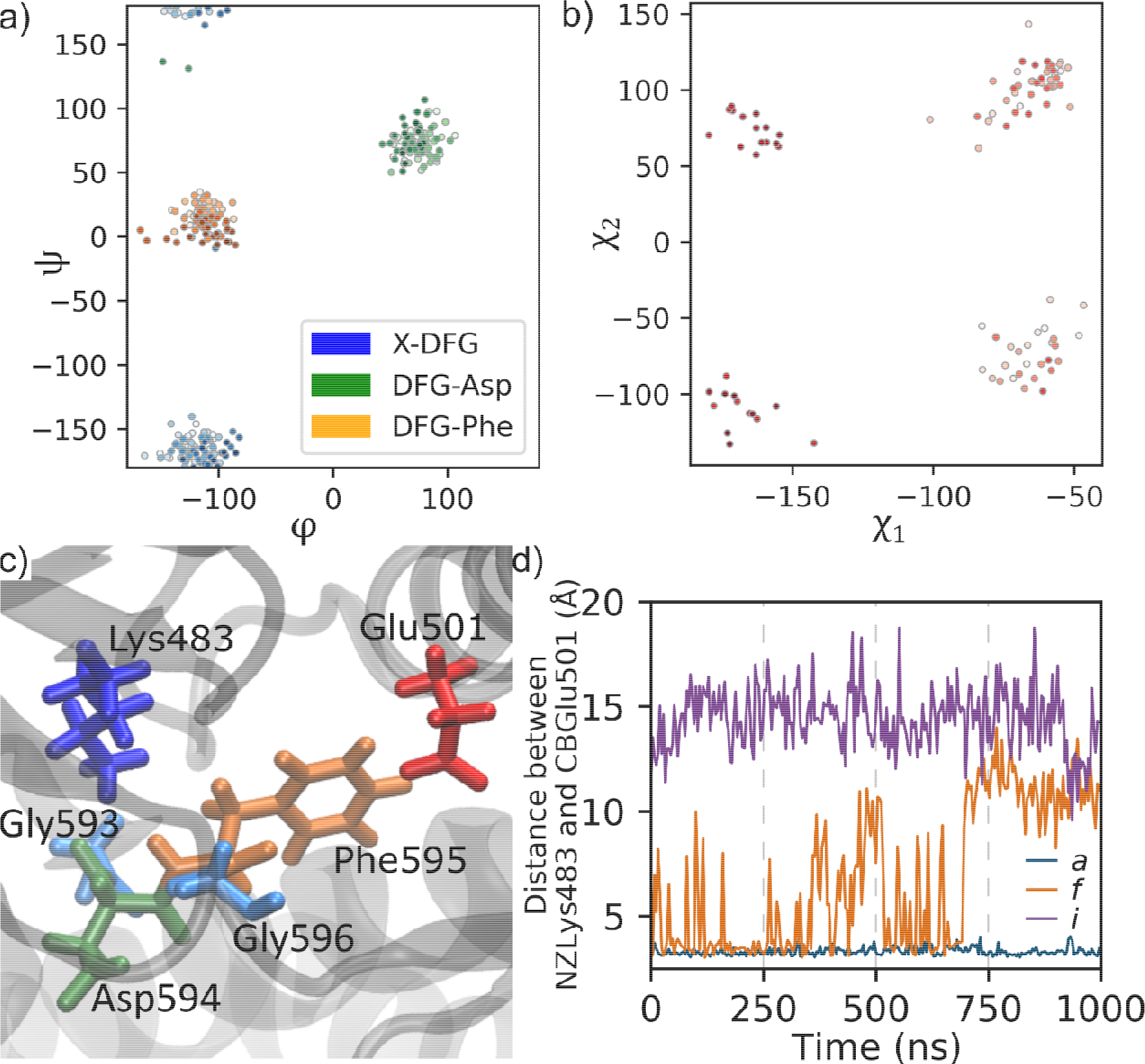
Phe595 trans χ_1_ rotamer disrupts the salt bridge between Lys483 and Glu501, resulting in an outward αC-helix. a) Ramachandran graph, b) sidechain dihedral angles, and c) representative snapshot of the residues ^593^GDFG^596^ for configuration *f*. Lightly colored symbols represent data from early in the simulation, darkly colored symbols are from the end of the simulation. d) Distance between the amide nitrogen of Lys483 and the carboxylate carbon of Glu501 for configuration *f* (orange). A representative system in which the αC-helix is always in (configuration *a*, blue) and always out (configuration *i*, purple) are provided for comparison.

For the remaining inactive B-Raf structures (configurations *g*-*i* for B-Raf V600E, *l*-*m* for wild-type B-Raf, *o*-*p* for pThr599/pSer602 B-Raf) we see two general trends. First, none of these inactive structures ever displayed the active BLAminus XDFG orientation. Second, the inactive wild-type B-Raf configurations and inactive B-Raf V600E configurations displayed different behavior around the DFG motif, even when starting from the same initial configuration. Configuration *g* (B-Raf V600E, intermediate αC-helix) started with a BLAplus orientation and changed to a BLAtrans orientation. The BLAtrans orientation matches the final orientation of configuration *f*, which was based on the same crystal structure as configuration *g*. Configuration *l* (wild-type B-Raf, intermediate αC-helix based on the same crystal structure as configurations *f* and *g*) also began with a BLAplus orientation (like configuration *g*), but ended with an ABAminus orientation. This marks a larger change away from the active configuration for the wild-type structure than the V600E structure. Configuration *o* (pThr599/pSer603 B-Raf, intermediate αC-helix based on the same crystal structure as configurations *f* and *g*) likewise changed from BLAplus to BLBplus. The B-Raf V600E mutant with an outward αC-helix (configuration *i*) and the phosphorylated B-Raf based on the same structure (configuration *p*) changed from BLBplus to BLBtrans and the mutant changed again to BLBminus, while the wild-type (configuration *m*) maintained the BLBplus throughout the simulation. Taken together, we see that in the inactive configurations, the mutant and wild-type structures based on the same initial PDBs do not exhibit the same dynamics around the DFG motif through the course of the simulations. This can also be seen in the cross correlation data in Figure S10: while the configurations with an inward orientation of the αC-helix all display similar results, in configurations with intermediate or outward αC-helix the results from the V600E mutant and pThr599/pSer602 match each other fairly well, while the wild-type simulations do not match each other, even when starting from the same PDB structures.

Another way to visualize the link between the αC-helix and the activation loop is through the radius of gyration (R_g_). Figure 11 shows the probability distribution of the radius of gyration versus the distance between Lys483 and Glu501. The general trend is that systems with an outward orientation of the αC-helix (large distance between Lys483 and Glu501) display a compact A-loop with low value of R_g_. Systems with an inward αC-helix with small distance between Lys483 and Glu501, conversely, have an extended A-loop with large values of R_g_. More specifically, these results show that the B-Raf V600E simulations sample more of the phase space as shown by the broader regions of high probability. Both the V600E and pThr599/pSer602 simulations that began with an inward αC-helix appear to have two distinct peaks in the probability distribution, while the wild-type has only one peak. For the intermediate and outward αC-helix, the pThr599/pSer602 and wild-type results behave similarly, while the B-Raf V600E simulations display different behavior. While the wild-type and phosphorylated B-Raf that began with an intermediate αC-helix relaxes to a state similar to the outward αC-helix simulations, the B-Raf V600E simulations with an intermediate αC-helix appear to occupy an intermediate state. This trend coupled with the XDFG dihedral results shows that the activation loop position has a large impact on the αC-helix.

**Figure 11.**
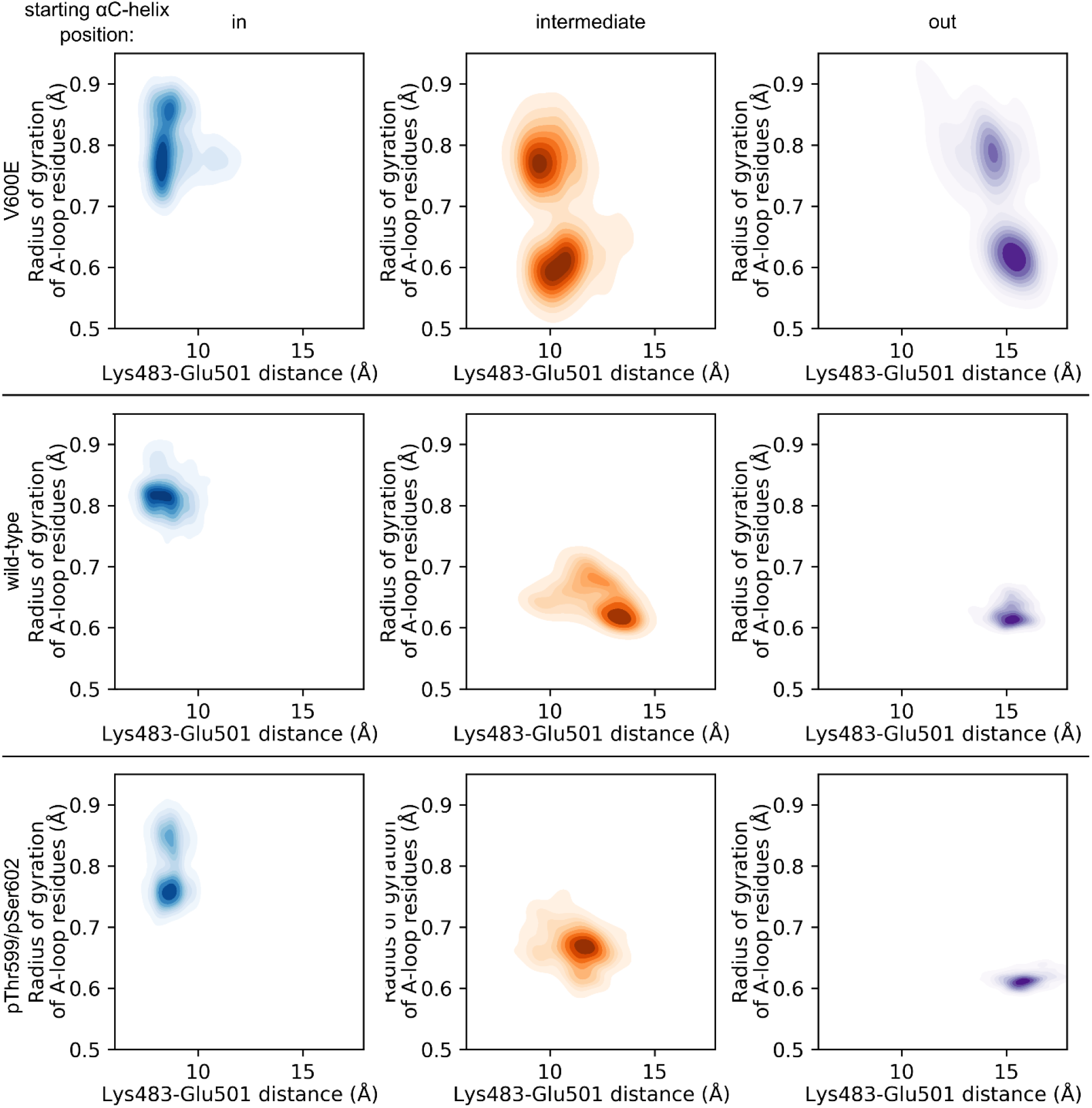
Probability distribution of radius of gyration of activation loop residues versus the distance between Lys483 and Glu501 shows that the B-Raf V600E or pThr599/pSer602 B-Raf result in a more flexible A-loop. Simulations that began with an inward orientated αC-helix are in blue, intermediate αC-helix in orange, and outward αC-helix in purple. Results from B-Raf V600E simulations are shown in the top row, wild-type simulations are in the middle row and pThr599/pSer602 simulations in the bottom row.

## 4 Discussion and Conclusion

B-Raf has been widely studied over the past decades.[2, 27, 89] Yet despite this, key features of B-Raf activation remain elusive. It is known that inactive B-Raf can exhibit, among other features, an outward placement of the αC-helix, an inhibitory loop called AS-H1 in the activation loop, and a disassembled regulatory spine.[39] In going from an inactive to the active conformation, B-Raf dimerizes with another Raf protein, has an inward placement of the αC-helix, an extended activation loop, and stacks four hydrophobic residues into a regulatory spine.[8] Understanding of how B-Raf moves between the active and inactive states could aid in targeted drug development. This is particularly important for the B-Raf V600E mutation, a so-called Class I mutation that adopts and active configuration independent of its upstream activator, Ras.

This work builds upon previous simulation studies of monomeric B-Raf to further our understanding of the mechanism of activation for B-Raf V600E. Previous MD simulations of monomeric B-Raf with an empty ATP binding pocket have found that wild-type B-Raf with an initially active configuration will quickly convert to an inactive conformation through the formation of the AS-H1 helix and an outward movement of the αC-helix.[85] We do not observe this in our simulations which include ATP and the magnesium ion, a requirement for a kinase action. Instead, active wild-type B-Raf remains primarily in the active configuration. The αC-helix predominately remains inward, as noted by the persistence of the Lys483-Glu501 salt bridge. The reason for these differences could be due to the interaction between the DFG-Asp residue and the magnesium ion in the ATP pocket. This interaction could stabilize the residues around the DFG motif, preventing the formation of the inhibitory helix. It has also been reported that active V600E will maintain an active conformation with an inward αC-helix.[85] While most of our simulations agree with this, there were results that indicate that the αC-helix of active B-Raf V600E can switch between the inward and outward conformations yet maintain an active BLAminus XDFG orientation. It is only when the DFG-Phe χ_1_ dihedral switched from minus to trans did the αC-helix become fixed in the outward position. This suggests that adopting the BLAminus XDFG orientation is a prerequisite for inward motion of the αC-helix. This insight could aid in the development of inhibitors that prevent the DFG motif from adopting this orientation.

Both B-Raf wild-type, phosphorylated wild-type, and V600E simulations which started with the AS-H1 helix maintained this inhibitory loop throughout the course of the simulation. Others have posited that the V600E mutation should disrupt this inhibitory loop due to the mutation of the hydrophobic Val with the negatively charged Glu. Although our simulations do not show the unwrapping of the inhibitory loop, the higher RMSF in the activation loop of the V600E simulations indicate that Glu mutation does increase the flexibility of this loop compared to Val. Similar results were seen in the phosphorylated wild-type simulations, which supports the hypothesis that the Val-to-Glu mutation mimics active loop phosphorylation, and the introduction of a negative charge to the AS-H1 loop is a key step towards activation. The persistence of the AS-H1 loop in our simulations, even with the Glu mutation, indicates that breaking this short helix could be the highest energy barrier in moving from the inactive to active configuration.

To conclude, we have performed extensive MD simulations on wild-type, pThr599/pSer603, and V600E B-Raf. For the first time these simulations include the physiologically important ATP molecule. Our results indicate that the V600E mutation results in activation via a mechanism that mirrors that seen in wild-type B-Raf. This has also been the case in PI3Kα and PTEN, and we expect is a general phenomenon.[90] We found that the αC-helix was only switched between the inward and outward orientation when the DFG motif adopted a BLAminus orientation and suggest that this information could assist in designing B-Raf inhibitors that exploit this feature.

## Author contributions

**Ryan Maloney**: Investigation, Formal Analysis, Writing – Original Draft. **Mingzhen Zhang**: Writing – Review & Editing. **Hyunbum Jang**: Conceptualization, Methodology, Writing – Review & Editing. **Ruth Nussinov**: Conceptualization, Supervision, Project administration, Funding acquisition, Writing – Review & Editing.

## Acknowledgements

This project has been funded in whole or in part with federal funds from the National Cancer Institute, National Institutes of Health, under contract HHSN26120080001E. The content of this publication does not necessarily reflect the views or policies of the Department of Health and Human Services, nor does mention of trade names, commercial products or organizations imply endorsement by the US Government. This research was supported [in part] by the Intramural Research Program of the NIH, National Cancer Institute, Center for Cancer Research. The calculations had been performed using the high-performance computational facilities of the Biowulf PC/Linux cluster at the National Institutes of Health, Bethesda, MD (https://hpc.nih.gov/).

