## Supplemental data for "The Mechanism of Activation of Monomeric B-Raf V600E"

| B-Raf region | Res ID | configuration | | | | | | | | | | | | | | | | |
| --- | --- | --- | --- | --- | --- | --- | --- | --- | --- | --- | --- | --- | --- | --- | --- | --- | --- | --- |
|  |  | 6U2G | a | b | c | d | e | f | g | h | i | j | k | l | m | n | o | p |
|  | Ile463 | X | X | X | X | X | X | X | X | X |  | X | X | X | X | X | X | X |
| P-loop | Gly464 | X | X | X |  | X | X | X |  | X |  | X | X |  | X | X | X | X |
|  | Ser465 | X | X | X | **HB** | X | **HB** | X |  | X |  |  |  |  | X | **HB** | X | X |
|  | Gly466 | X | X | X | X | X | **HB** | X | X | X | X | X | X | X | X | X | X | **HB** |
|  | Ser467 | **HB** | **HB** | **HB** | **HB** | **HB** | **HB** | **HB** | **HB** | X | X | **HB** | **HB** | **HB** | **HB** | **HB** | **HB** | **HB** |
|  | Phe468 | X | **HB** | **HB** | X | X | X | X |  | **HB** | **HB** | X |  |  | **HB** |  |  | **HB** |
|  | Gly469 | X | **HB** | **HB** |  |  |  |  |  |  | X |  |  |  | **HB** |  |  | **HB** |
|  | Val471 | X | X | **X** | X | X | X | X | X | X | X | X | X | X | X | X | X | X |
| B.P. | Ala481 | X | X | X | X | X | X | X | X | X | X | X | X | X | X | X | X | X |
|  | Lys483 | X | X | X | X | **HB** | **HB** | X | X | **HB** | **HB** | X | X | **HB** | **HB** |  | X | **HB** |
| B.P. | Leu514 | X | X | X | X | X | X | X | X | X | X | X | X | X | X | X | X | X |
| B.P. | Thr529 | X | **HB** | X | X | **HB** | X | **HB** | **HB** | X | X | X | **HB** | X | X | X | **HB** | X |
|  | Gln530 | **HB** |  | **HB** | **HB** | **HB** | X | **HB** | **HB** | **HB** | **HB** | **HB** | **HB** | **HB** | **HB** | **HB** | **HB** | **HB** |
| B.P. | Trp531 | X | X | **HB** | X | X | **HB** | X | X | X | X | X | X | X | X | X | X | X |
| B.P. | Cys532 | X |  | X | **HB** | **HB** | **HB** | X | X | **HB** | **HB** | **HB** | **HB** | **HB** | **HB** | **HB** | **HB** | **HB** |
| Hinge | Ser535 |  |  |  | X |  |  |  |  |  |  |  |  |  |  |  |  |  |
|  | Ser536 | X | X | X | X |  | X |  |  |  |  |  | X |  |  | X | X | X |
|  | His539 |  |  |  |  |  |  |  |  |  |  |  |  |  |  |  |  | X |
| Catalytic loop | Asp576 | **HB** | X | X | X |  |  | X |  |  | X | X |  |  | X | X |  | X |
|  | Lys578 | **HB** | X | X | X |  | X | X | X | **HB** | **HB** | X | X | X | X | **HB** | X | **HB** |
|  | Asn580 | X | **HB** | X | **HB** | **HB** | **HB** | X | X |  | X | X | X | X |  | **HB** | **HB** | X |
|  | Asn581 | **HB** |  | **HB** |  |  | X |  | X | X | X |  | X | X | **HB** | X | **HB** | X |
| C-spine | Phe583 | X | X | X | X | X | X | X | X | X | X | X | X | X | X | X | X | X |
|  | Gly593 |  | X |  |  |  |  |  |  |  | X |  |  |  |  |  |  |  |
| DFG | Asp594 | X | X | X | X | X | X | X | X | X | X | X | X | X | X | X | X | X |
|  | Gly596 |  |  |  |  | X |  |  |  |  |  |  |  |  |  |  |  |  |
| A-loop | Leu597 |  |  |  |  |  |  |  |  |  |  | X |  |  |  |  |  |  |
|  | Glu600 |  |  |  |  |  |  |  |  |  | X |  |  |  |  |  |  |  |
|  | Lys601 |  |  |  |  |  |  |  |  |  | **X** |  |  |  |  |  |  |  |
|  | Ser614 |  |  |  |  |  |  |  |  |  |  |  |  | X |  |  | X | X |
|  | Gly615 |  |  |  |  |  |  |  |  |  | X |  |  |  | X |  | X | X |
|  | Ser616 |  | X | X |  |  |  |  | X | X |  | X |  |  |  |  |  | X |

**Table S1** Summary of binding between B-Raf and ATP. “X” in the table indicates which residues are within 4 Å of ATP in the initial configuration. “HB” indicates the residues that form hydrogen bonds with ATP. A blue background indicates that a salt bridge between ATP and the corresponding residue. A green background indicates π-π stacking between that residue and the adenine ring of ATP. “B.P.” indicates the ATP binding pocket as defined in Tsai *et al*. (PNAS 2008, 105:3041). Binding between 6U2G and the ATP analog AMP-PCP is provided for comparison.

| PDB ID | PDB title | chain used | Mutated residues | Missing residues |
| --- | --- | --- | --- | --- |
| 3OG7 | B-Raf Kinase V600E oncogenic mutant in complex with PLX4032 | A | Lys522Ala  Ile543Ala  Ile544Ser  Ile551Lys  Gln562Arg  Leu588N  **Val600Glu**  Lys630Ser  Phe667Glu  Trp673Ser  Ala688Arg  Leu706Ser  Gln709Arg  Ser713Glu  Leu716Glu  Ser720Glu | 545-547  597-614  627-630 |
| 4MNE | Crystal structure of the BRAF:MEK1 complex | B | none | 465-468 |
| 4MNF | Crystal structure of BRAF-V600E bound to GDC0879 | A | **Val600Glu** | 601-615 |
| 4XV2 | B-Raf Kinase V600E oncogenic mutant in complex with Dabrafenib | A | Ile551Lys  Gln562Arg  Leu588Asn  **Val600Glu**  Lys630Ser  Phe667Glu  Tyr673Ser  Ala688Arg | 488-489  597-614  627-631 |
| 5ITA | Crystal structure of BRAF kinase domain bound to AZ-VEM | A | Ile551Lys  Gln562Arg  Leu588Asn  Lys630Ser  Phe667Glu  Tyr673Ser  Ala688Arg  Leu706Ser  Gln709Arg  Ser713Glu  Leu716Glu  Ser720Glu | 598-615  629-630  719-720 |
| 6U2G | BRAF-MEK complex with AMP-PCP bound to BRAF | B | none | 603-610 |
| 6U2H | BRAF dimer bound to 14-3-3 | D | none | 608-610 |

**Table S2** Summary of PDB entries used to create the initial configurations for simulations.

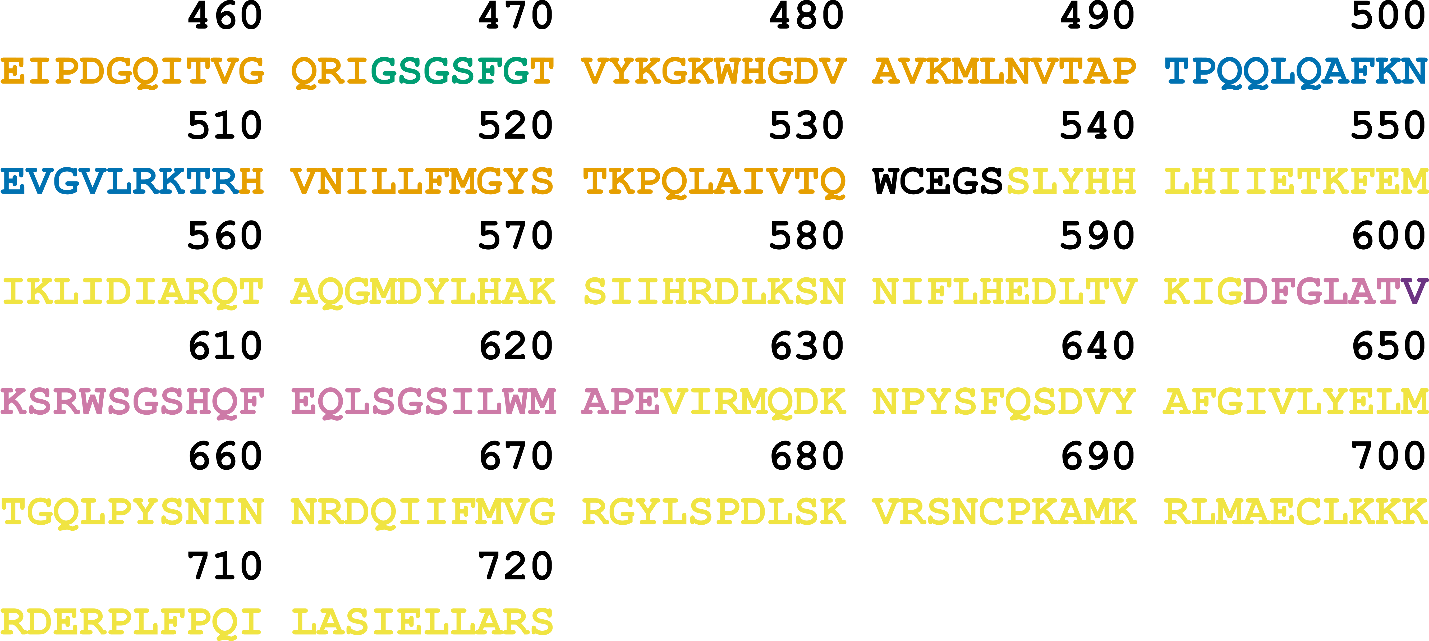

**Figure S1** B-Raf kinase domain residues 451 to 720. Residues are colored to match Figure 1b: N-lobe (orange), P-loop (green), αC-helix (blue), hinge (black) C-lobe (yellow), and A-loop (light purple). The Val600 residue is colored dark purple.

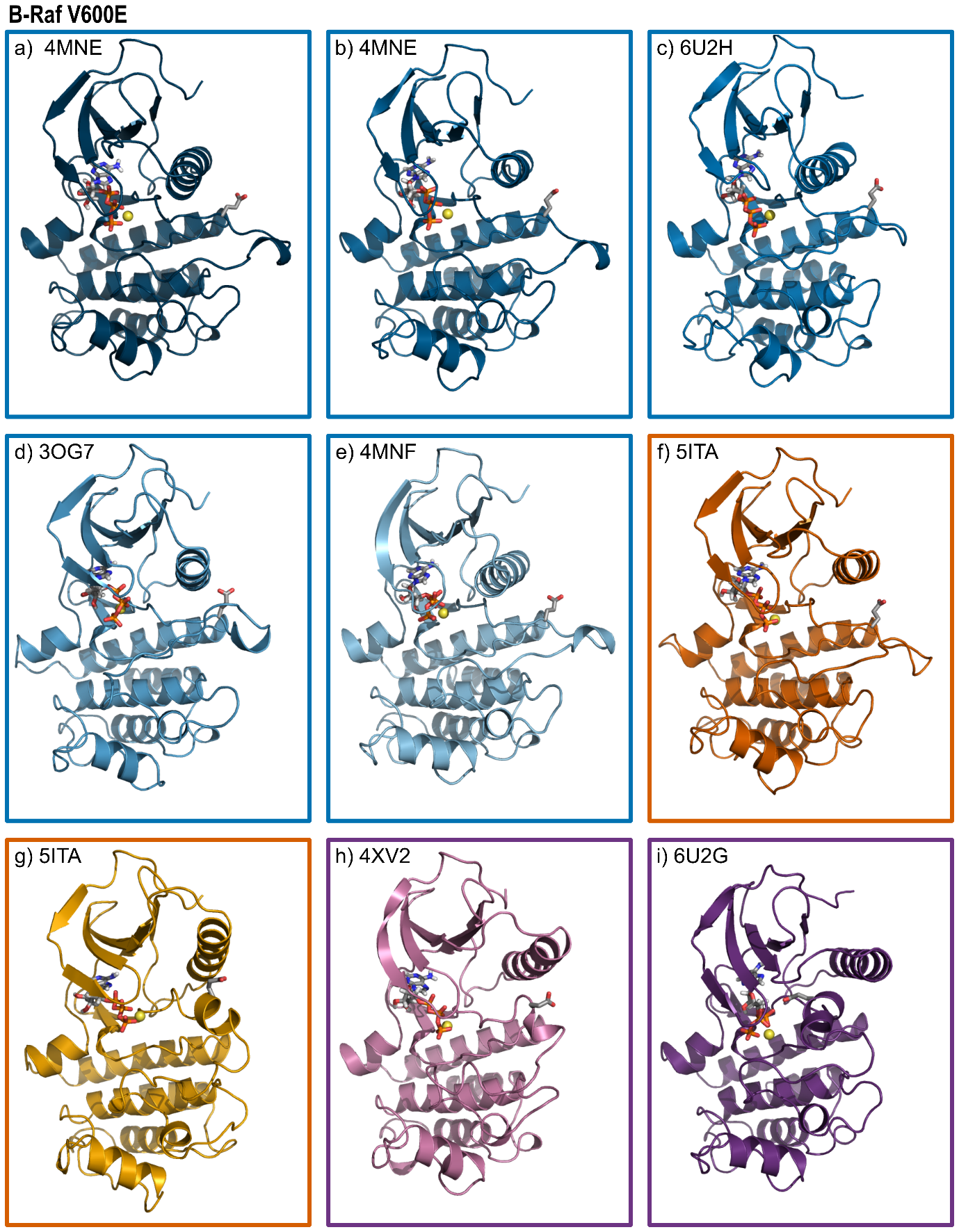

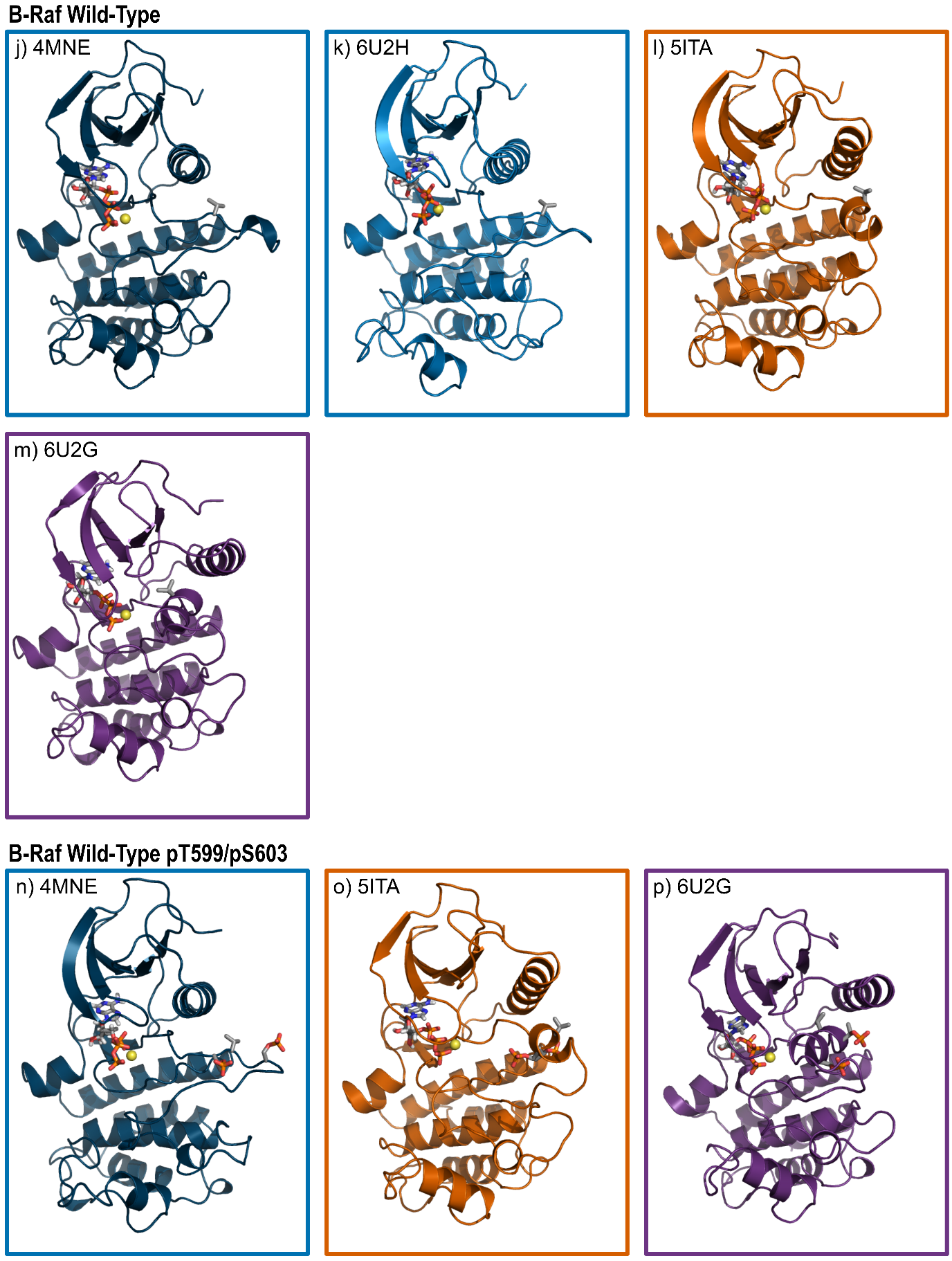
**Figure S2** Initial configurations for B-Raf V600E (*a*-*i*), wild type (*j*-*m*), and wild type with pThr599/pSer603 (*n*-*p*) simulations. The color of the box around each snapshot indicates the initial position of the αC-helix (inward blue, intermediate orange, outward purple).

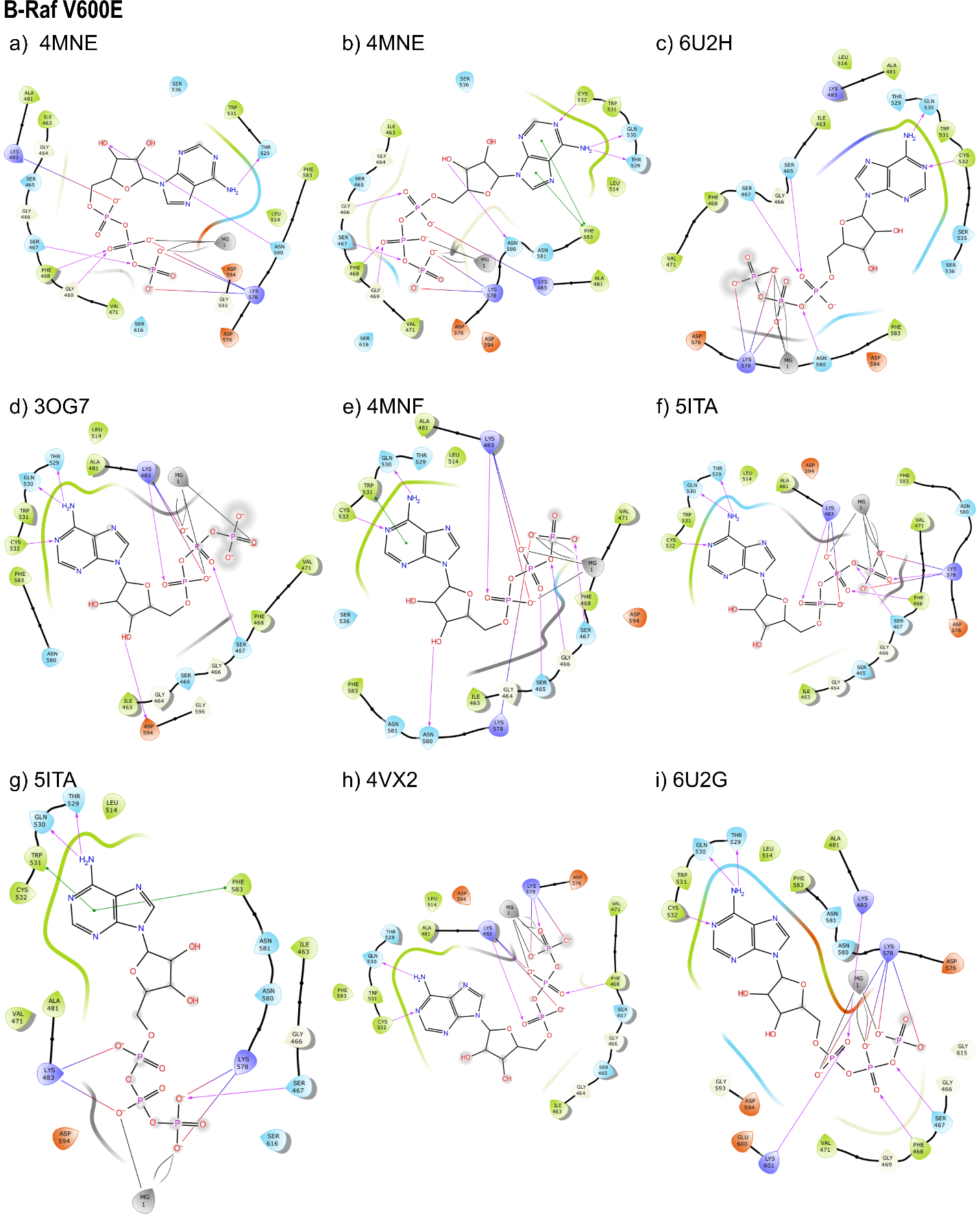

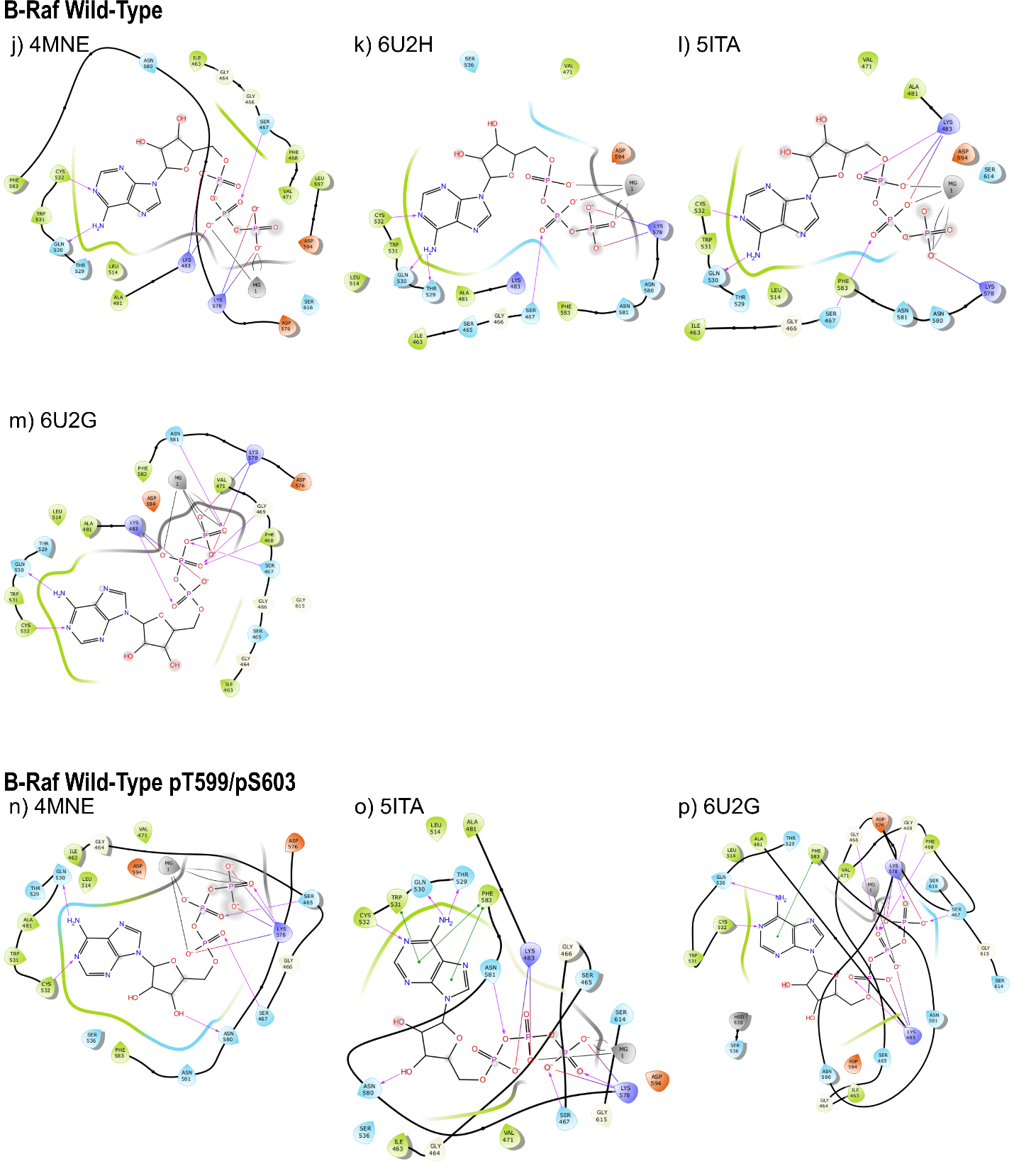

**Figure S3** Ligand interaction diagrams between ATP and B-Raf.

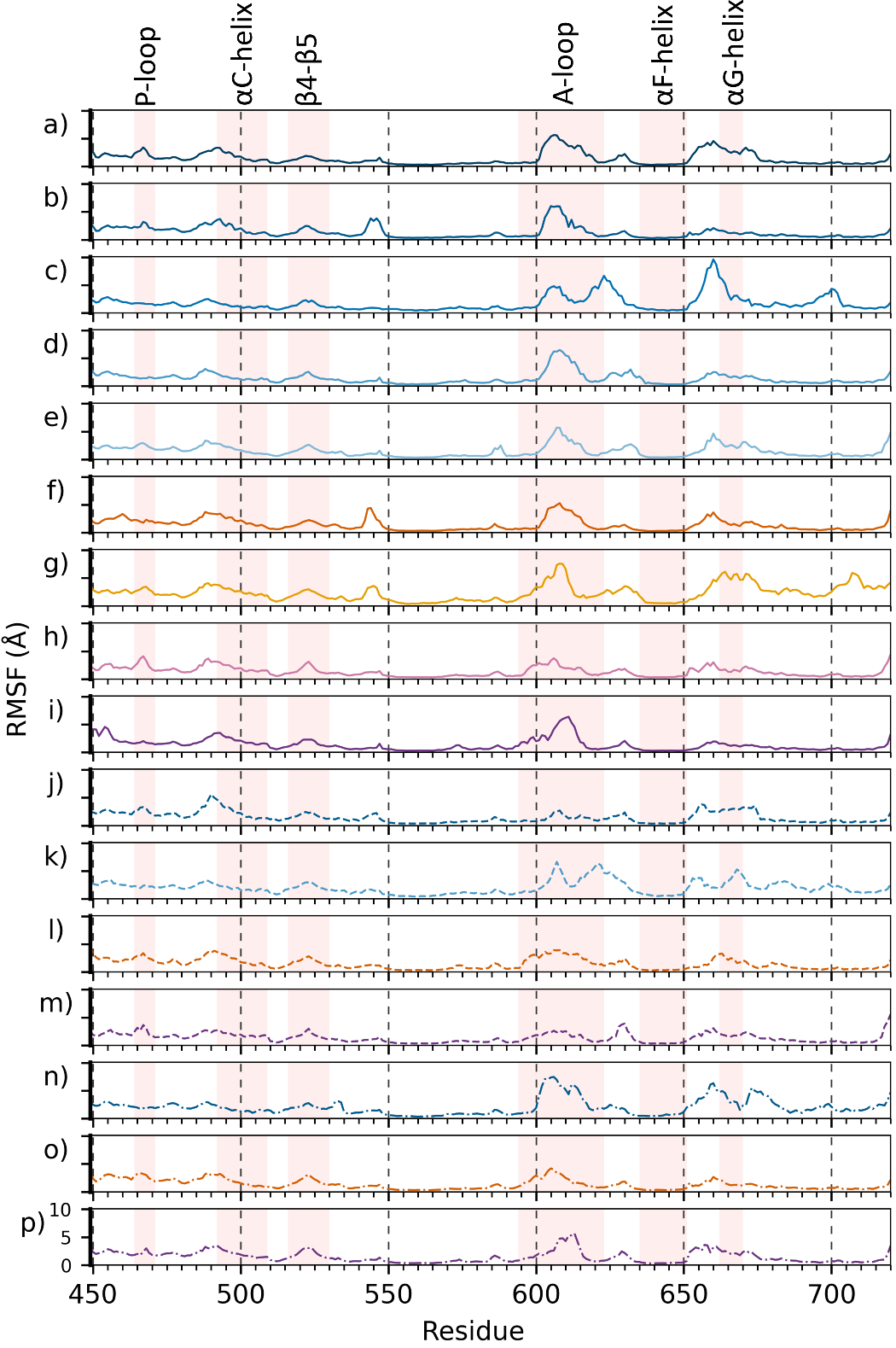
**Figure S4** Root mean square fluctuation (RMSF) versus residue number for each of the simulated configurations. B-Raf V600E simulation results (*a*-*i*, solid line), B-Raf wild type simulation results (*j*-*m*, dashed line), and pThr599/pSer602 B-Raf (*n*-*p*, dash-dot line). The y-axis spans the same range in all plots and is indicated in *p*. The line color indicates the initial position of the αC-helix (inward blue, intermediate orange, outward purple).

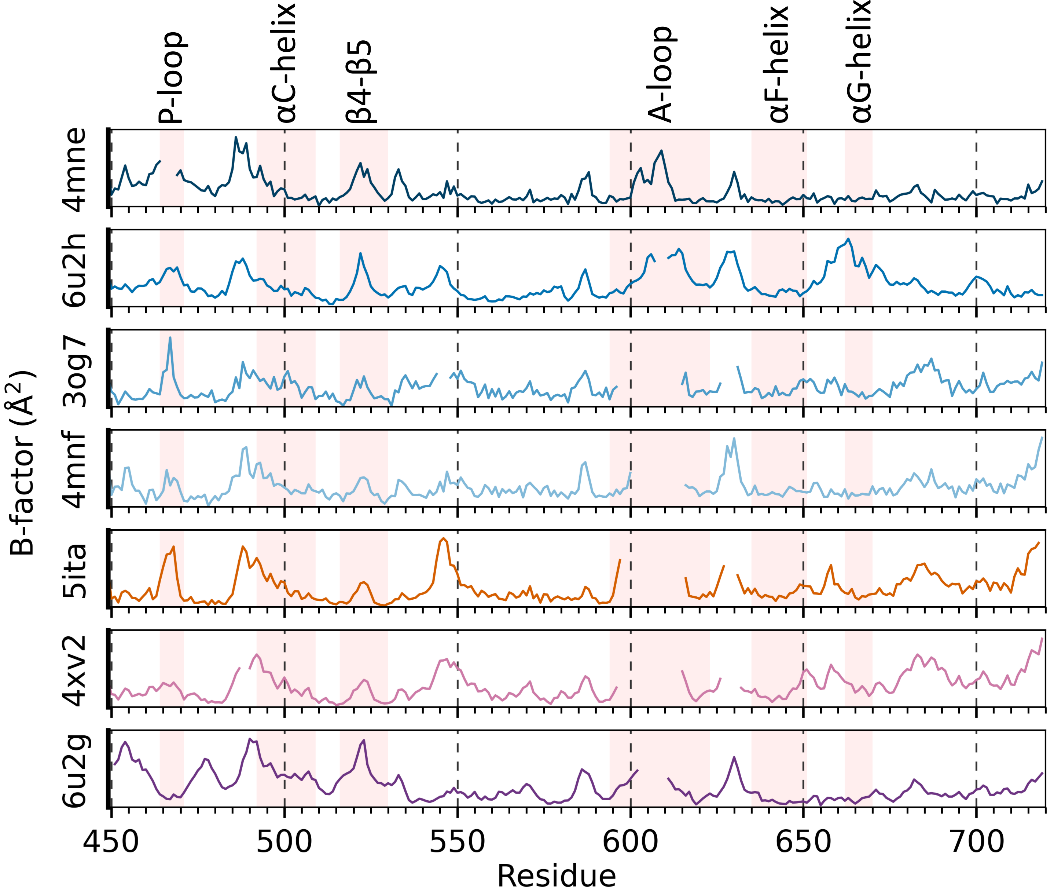
**Figure S5** *B*-Factors for PDB versus residue number for each of the crystal structures used to construct the initial configurations. The line color indicates the position of the α-C helix (inward blue, intermediate orange, outward purple).

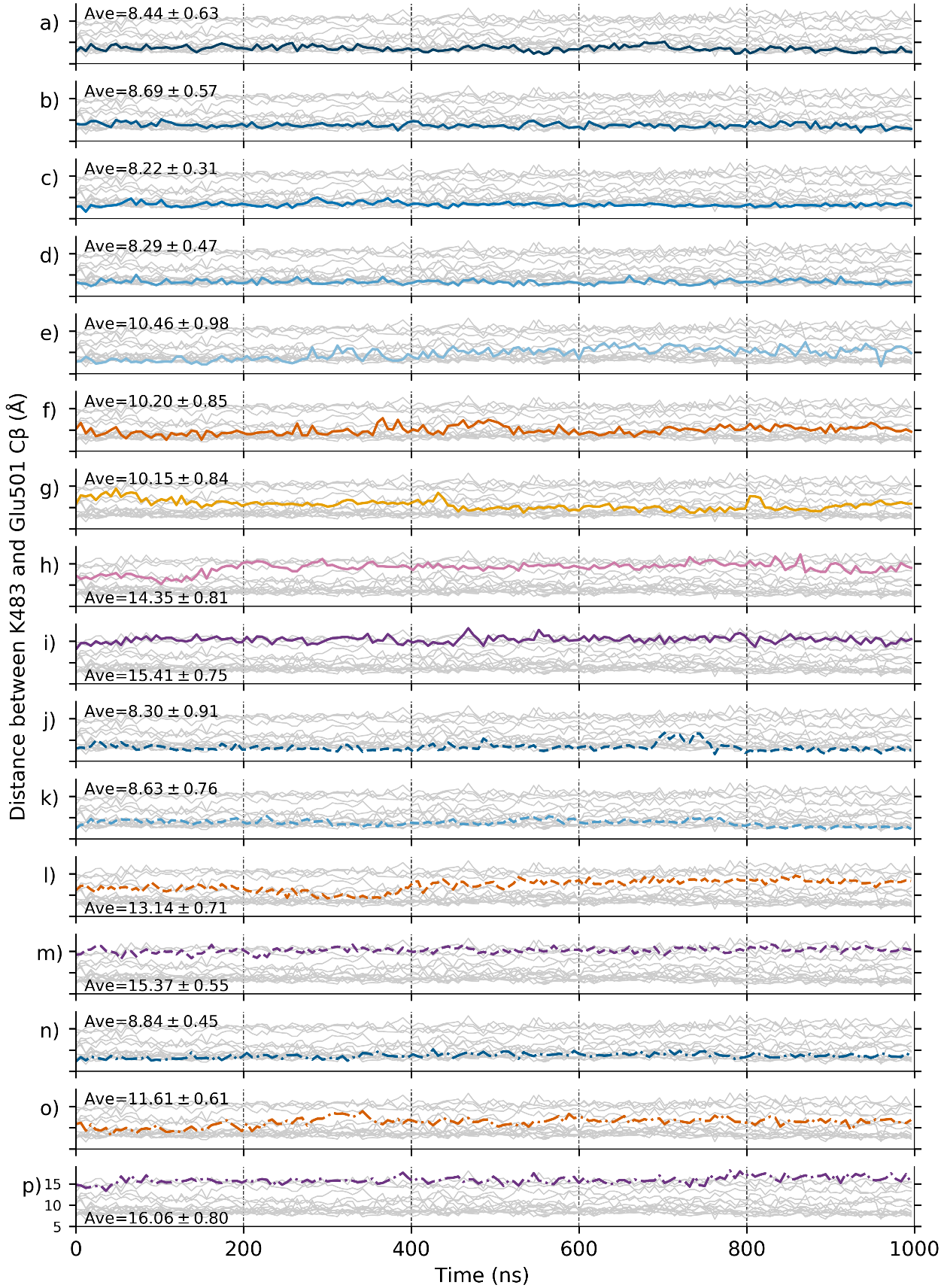
**Figure S6** Distance between CB on Lys483 and CB on Glu501 over the course of the simulation. B-Raf V600E simulation results (a-i, solid line), B-Raf wild type simulation results (j-m, dashed line), and pThr599/pSer602 B-Raf (n-p, dash-dot line). The y-axis spans the same range in all plots and is indicated in p. Line color indicates the initial position of the αC-helix (inward blue, intermediate orange, outward purple). The gray lines in the background depict results from the other simulations for comparison. Average values taken over final 600 ns of the simulation.

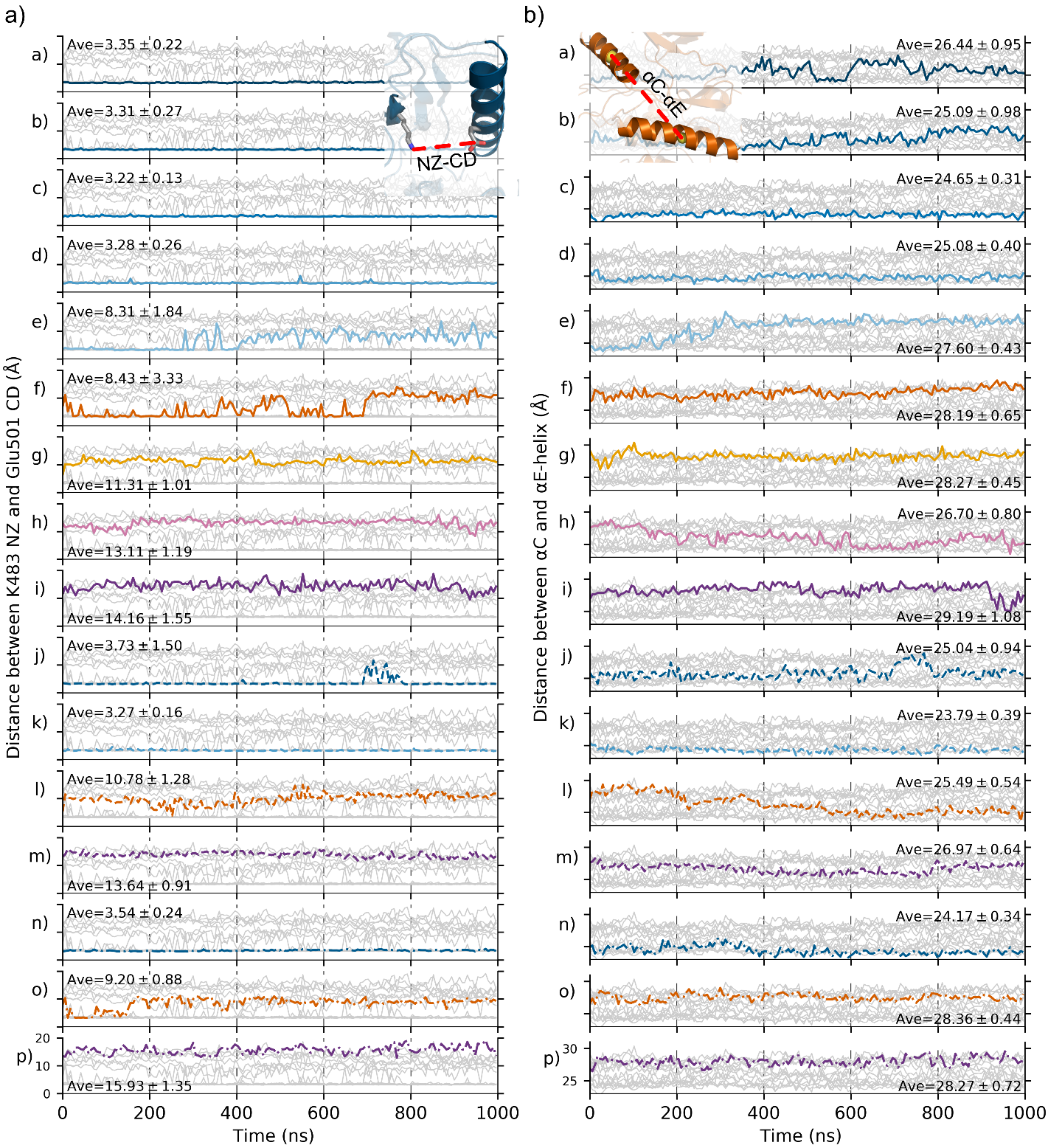
**Figure S7** a) Distance between the atom pair NZ on Lys483 and CD on Glu501 over the course of the simulation. b) Center of mass distance between the αC-helix in the N-lobe and αE-helix in the C-lobe over the course of the simulation. B-Raf V600E simulation results (a-i, solid line), B-Raf wild type simulation results (j-m, dashed line), and pThr599/pSer602 B-Raf (n-p, dash-dot line). The y-axis spans the same range in all plots and is indicated in p. Line color indicates the initial position of the α-C helix (inward blue, intermediate orange, outward purple). The gray lines in the background depict results from the other simulations for comparison. Average values taken over final 600 ns of the simulation.

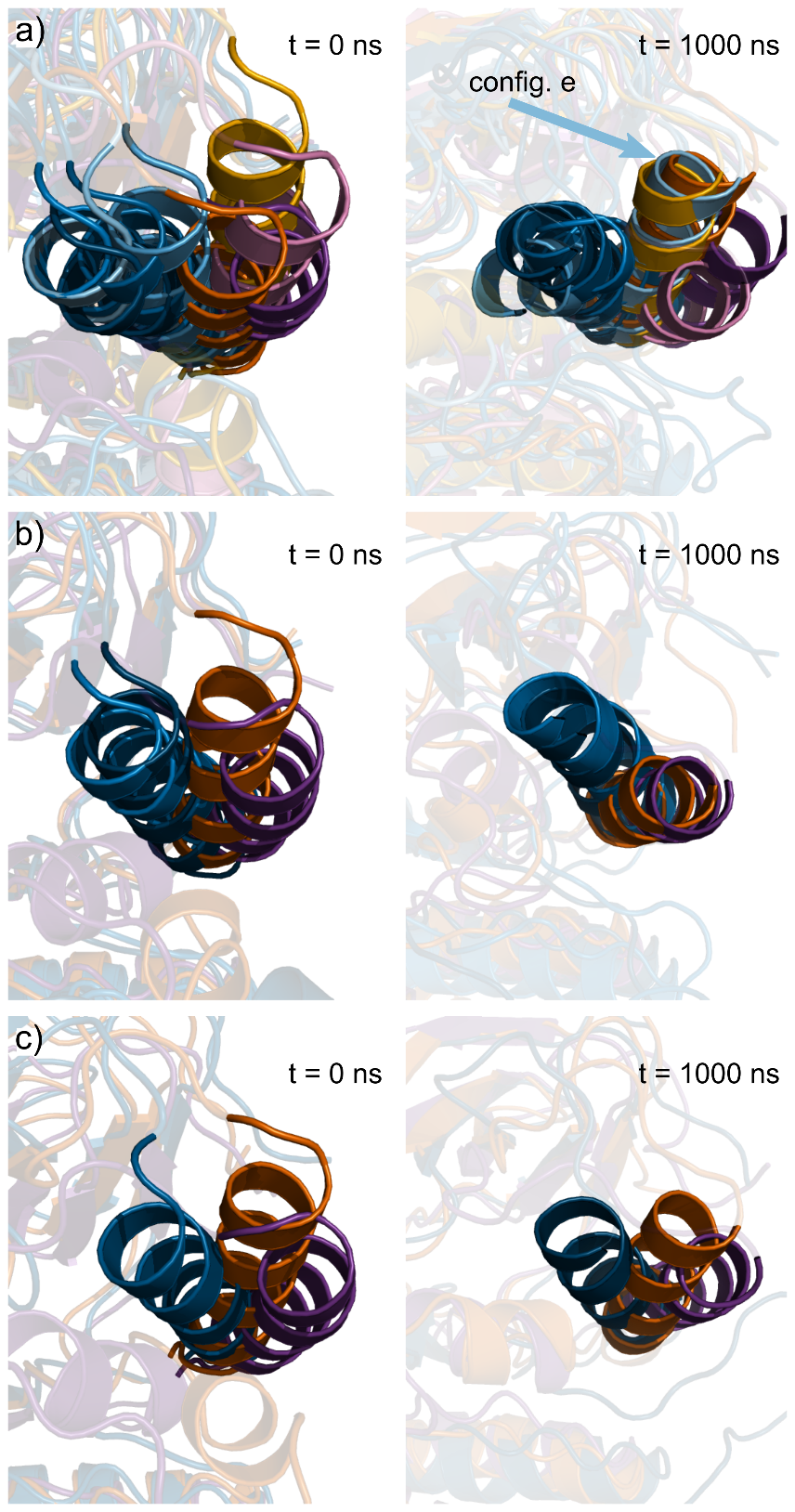
**Figure S8** Initial and final configurations of the αC-helix for a) B-Raf V600E, b) wild-type B-Raf, and c) pThr599/pSer602. Simulations that began with an initial inward αC-helix are shown in blue, intermediate αC-helix are shown in orange, and outward αC-helix shown in purple.

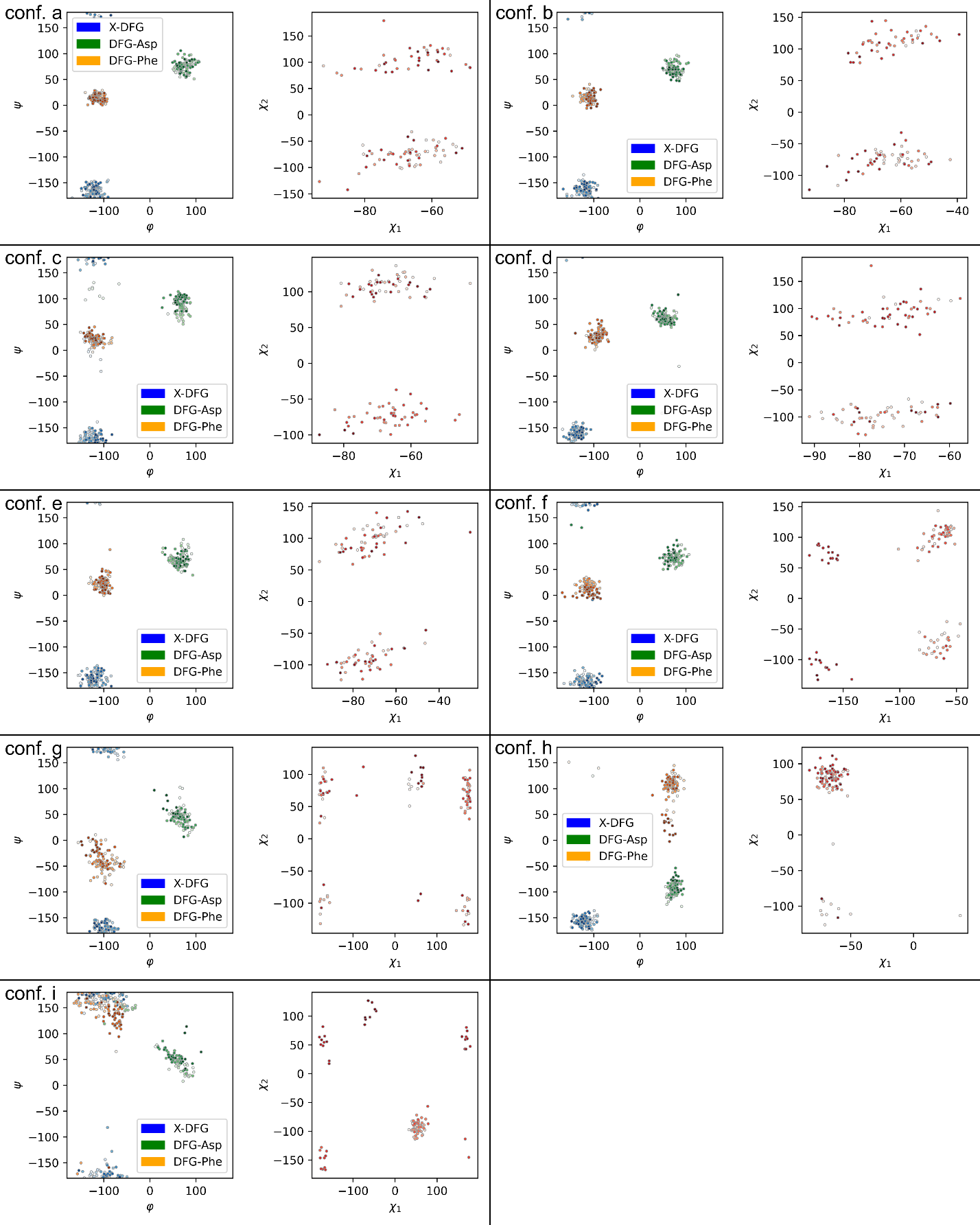

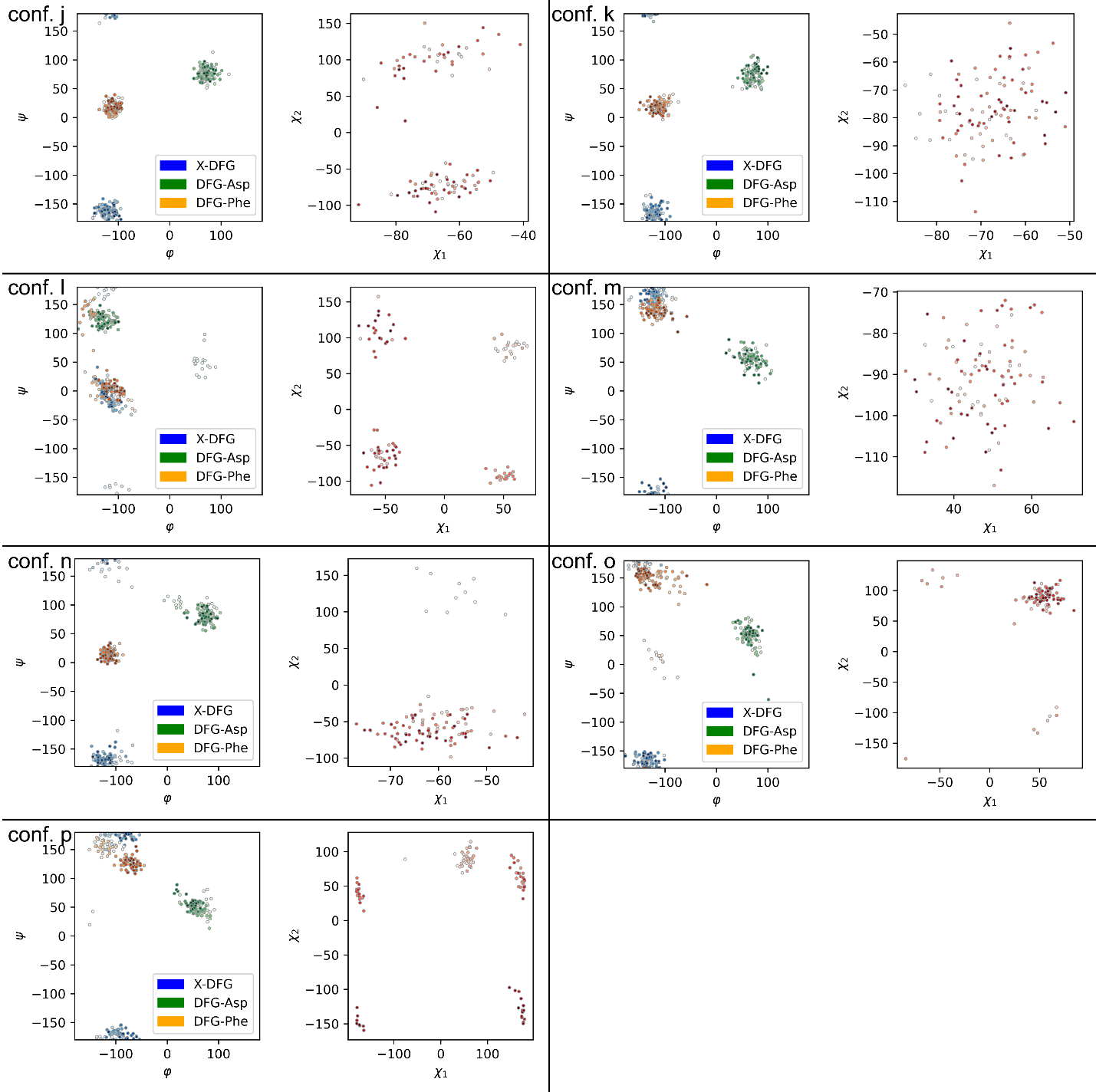

**Figure S9** Ramachandran graphs and sidechain dihedral angles of the residues ^593^GDFG^596^ for B-Raf V600E (conf. *a*-*i*), B-Raf wild type (conf. *j*-*m*) and pThr599/pSer602 B-Raf (conf. *n*-*p*).

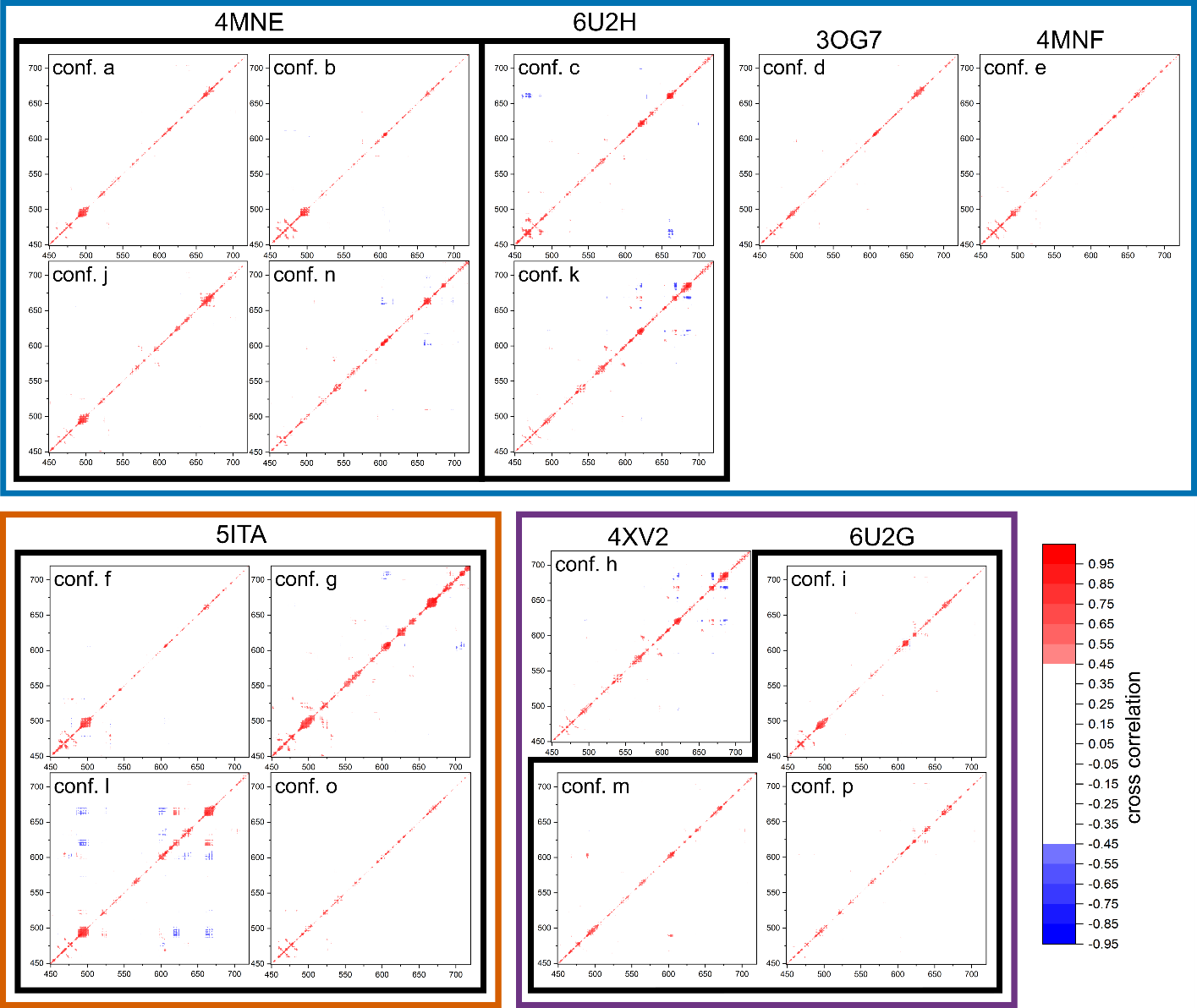
**Figure S10** Cross correlation of B-Raf kinase domain residues 449–720 for simulations that begin with an inward αC-helix (blue box) intermediate αC-helix (orange box) or outward αC-helix (purple box). Black boxes indicate configurations based on the same initial crystal structure. Configurations a-i are simulations of B-Raf V600E, configurations j-m are wild-type B-Raf, and configurations n-p are pThr599/pSer602 B-Raf.
